# Mechanostimulatory cues determine intestinal fibroblast fate and profibrotic remodeling in a physiodynamic human gut-on-a-chip

**DOI:** 10.1101/2025.06.07.658434

**Authors:** Soyoun Min, Nam Than, Yong Cheol Shin, Elif G. Ertugral, Chandrasekhar R. Kothapalli, Olumuyiwa Awoniyi, Hyun Jung Kim

## Abstract

Biomechanical cues, including shear stress and mechanical strain, are key regulators of intestinal cellular behavior, yet their mechanostimulatory impact on fibroblasts responses during early fibrotic remodeling remains poorly understood. Using a bioengineered gut-on-a-chip model, we independently modulated flow and mechanical strain to assess fibroblast dynamics under intact or impaired epithelial barriers. Inflammation-associated fibroblasts resisted biomechanical stress, exhibiting myofibroblast-like phenotypes with hypertrophy and elevated α-smooth muscle actin aligned with stress fibers. In contrast, normal fibroblasts were highly susceptible to shear stress, undergoing matrix metalloproteinase-dependent apoptotic injury, while mechanical strain alone had minimal effect. Notably, an intact epithelial barrier was both necessary and sufficient to protect fibroblasts from shear-induced damage, suggesting that “good fences make good neighbors”. Under barrier dysfunction, prolonged exposure to shear stress induced the formation of stiff fibroblast aggregates composed of mechanoadaptive myofibroblast-like cells. These findings identify mechanostimulatory cues, particularly shear stress, as critical drivers of early fibrotic remodeling in inflammatory bowel disease and underscore epithelial barrier integrity as an essential biomechanical safeguard against pathological fibroblast dysregulation.

## Introduction

Intestinal fibrosis in inflammatory bowel disease (IBD) results from recurrent and localized inflammation, leading to tissue thickening, scarring, strictures, and stenosis^1–3^. In both Crohn’s disease (CD) and ulcerative colitis (UC), mucosal fibrosis is linked to chronic inflammation and epithelial barrier dysfunction, exposing underlying fibroblasts to abnormal luminal fluid shear and disrupting extracellular matrix (ECM) remodeling, ultimately impairing bowel function^4^. Despite its high prevalence, no targeted therapies exist for IBD-associated fibrosis, as its primary drivers remain unclear^5^. Notably, a subset of IBD patients, prevalently in CD, develops severe fibrosis^6–10^ and exhibits poor responsiveness to biologic therapies, including anti-tumor necrosis factors (TNF) blockers^11, 12^, indicating that fibrosis progression is not solely driven by inflammation. Hence, identifying key profibrotic triggers beyond inflammation is essential to advance our understanding of early fibrogenic processes and develop effective anti-fibrotic therapies.

The intestinal mucosa is inherently subjected to rhythmic mechanical deformations and luminal shear stress during peristalsis, which are essential for maintaining intestinal homeostasis^13^. In IBD, chronic inflammation often disrupts the epithelial barrier, creating damaged mucosal loci that serve as initiation sites for early fibrotic remodeling^14^. Barrier dysfunction exacerbates biomechanical stress on subepithelial fibroblasts, exposing them to abnormal physical forces. However, the extent to which these forces drive fibroblast activation, excessive ECM deposition^15^, secretion of fibrogenic cytokines^16^, responsiveness to microbial signals^17^, and differentiation into myofibroblasts^18^ remains poorly understood. Furthermore, dysregulated intestinal peristalsis in IBD further amplifies mechanobiological stress on stromal cells, often associated with abnormalities in intestinal smooth muscle cells (SMCs)^19–23^. Despite these associations, the specific contribution of luminal shear stress and mechanical deformation to early fibrotic cascades remain unclear. Investigating the independent role of pathobiological and biomechanical factors has been hindered by the confounding complexity of *in vivo* systems. Hence, a modular intestine model capable of recapitulating the mucosal microenvironment and mechanobiological interactions is essential to elucidate the mechanostimulatory cues governing fibroblast activation and profibrotic remodeling in IBD.

We have developed a bioengineered human gut-on-a-chip microphysiological system (MPS) that can demonstrate three-dimensional (3D) morphogenesis of intestinal epithelial cells^24–27^, peristalsis-like physiodynamic motions and fluid shear stress^28–30^, multi-cellular host-microbiome co-cultures including anaerobic gut bacteria^28, 31^, pathomimetic modeling of human intestinal disorders^32–34^, and integrative cultures with human intestinal organoids for conducting precision medicine studies. This modular system facilitates the identification of specific pathophysiological factors by selectively introducing or removing target cell types or contributing components in a controlled spatiotemporal manner^25, 33^. For example, repetitive deformations in a gut-on-a-chip emulate the mechanical contribution of SMCs in the human intestine. This approach exclusively isolates the mechanobiological effects of SMCs on intestinal mucosal cells, eliminating confounding biological factors and reducing *in vivo* complexity for mechanistic studies. Additionally, the gut-on-a-chip enables independent control of fluid flow and cyclic stretching, providing precise assessment of the mechanomodulatory milieu affecting mucosal cells. Notably, replicating these biomechanical cues in animal models remains a significant challenge.

In this study, we leveraged a physiodynamic gut-on-a-chip model to investigate the role of biomechanical cues in early fibrosis process. By independently controlling fluid flow and mechanical deformation, we identified how these mechanostimulatory factors can drive fibroblast reprogramming and trigger early profibrotic remodeling. We also elucidated the essential role of a protective epithelial barrier in regulating mechanodynamic stress during the initial stages of fibrosis in IBD. Our findings highlight the potential of a modular gut-on-a-chip platform to recapitulate profibrotic microenvironments, providing critical insights into the mechanobiological drivers of fibrosis, and identifying potential therapeutic targets.

## Results

### Inflammation-associated fibroblasts retain disease-specific cellular characteristics

To investigate the mechanostimulatory responses of intestinal fibroblasts associated with profibrotic process in IBD, we examined the morphological and functional characteristics of normal fibroblasts (nFib) and inflammation-associated fibroblasts (iFib) isolated from a colonic biopsy of a UC patient with severe inflammatory injury. When these cells were cultured on a rigid surface of a standard T75 flask (stiffness at 3.07±0.18 GPa, Supplementary Fig. 1a), nFib exhibited random orientations and typical fibroblast morphology, whereas iFib displayed a striped, elongated appearance (Fig. 1a, Supplementary Fig. 1b). At confluence, nFib maintained a stochastic orientation, while iFib demonstrated anisotropic alignment with a defined orientation index (Fig. 1b, Supplementary Fig. 1b). Quantitative analysis of cell shapes revealed that iFib had a significantly higher aspect ratio (4.20±0.41, *p*<0.001) compared to nFib (1.78±0.17), confirming distinct morphological differences between the two fibroblast types (Fig. 1c). Furthermore, molecular profiling of nFib and iFib cultured on T75 flasks confirmed that iFib exhibited elevated expression of α-smooth muscle actin (α-SMA) and fibronectin among CD90+ and vimentin+ cells, as quantified by flow cytometry (Fig. 1d, Supplementary Fig. 1c). These findings confirmed the distinct morphological and molecular characteristics of inflammation-associated fibroblasts compared to normal fibroblasts.

**Fig. 1.**
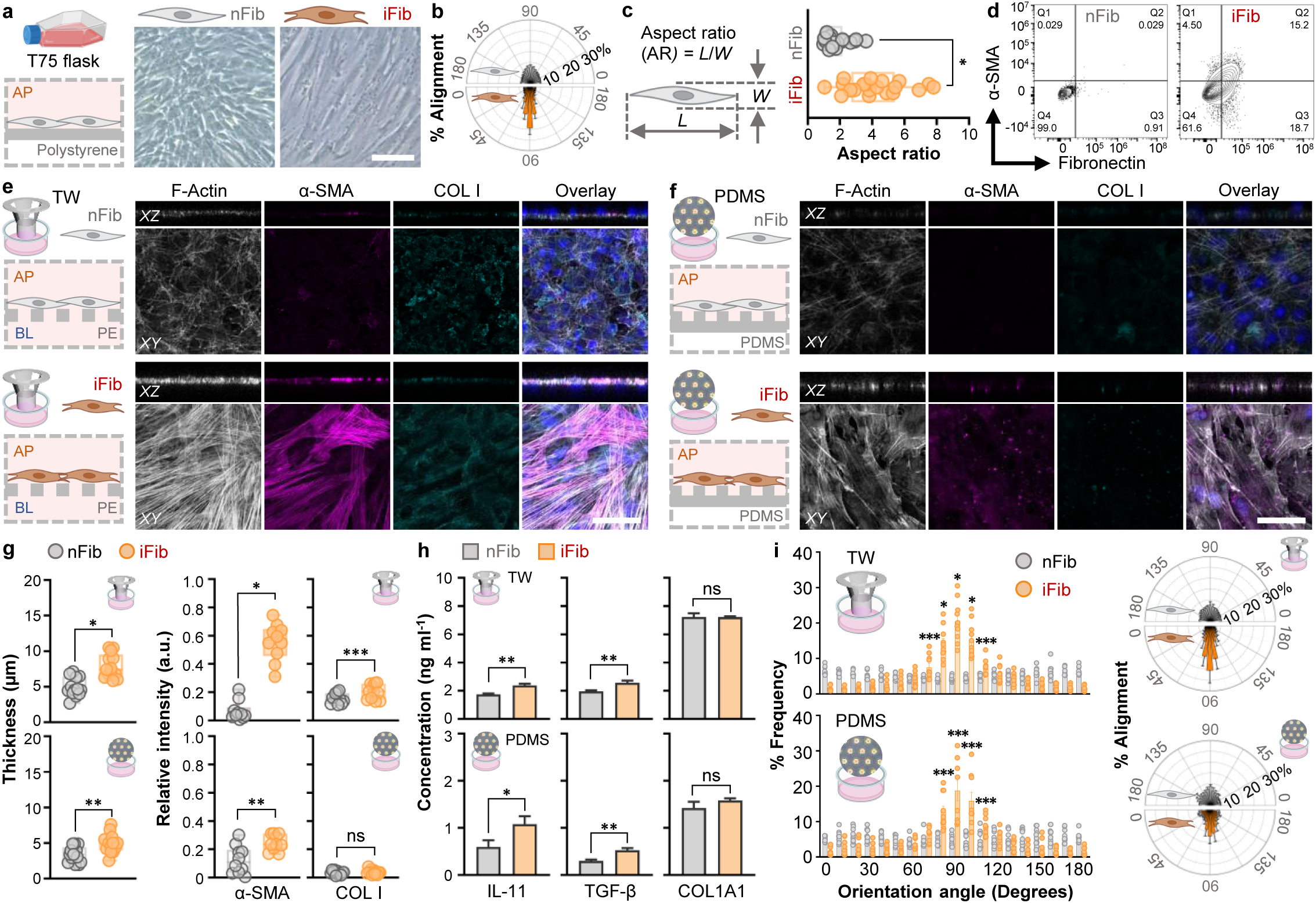
Inflammation-associated fibroblasts retain profibrotic characteristics influenced by substrate stiffness and culture conditions. **a**, Phase-contrast micrographs of normal fibroblasts (nFib) and inflammation-associated fibroblasts (iFib) cultured on rigid T75 flask surfaces for 110 h. **b**, Orientation index of nFib and iFib cells, quantitatively assessed as a function of probability alignment, demonstrating cellular orientation on T75 flasks after 110 h of cultures (*n*=3). **c,** Aspect ratio profiles of nFib and iFib cells cultured on T75 flasks for 110 h (*n*=20). **d**, Flow cytometric analysis of nFib and iFib cultured on T75 flasks, highlighting fibrotic markers α-SMA and fibronectin. **e**, Morphological and functional characteristics of nFib and iFib cultured on a Transwell (TW) with polyester nanoporous inserts (pore size: 0.4 µm) for 5 days. Immunofluorescence micrographs show top-down (*XY*) and cross-sectional (*XZ*) views with markers for F-actin (grey), α-SMA (magenta), COL I (cyan), and their overlay with nuclei (blue) (*n*=5). **f**, Morphological and functional characteristics of nFib and iFib cultured on PDMS membranes (10:1 w/w, prepolymer: curing agent) on glass-bottom well plates for 5 days, visualized under the same immunofluorescence conditions as in panel **e**. **g**, Quantification of cell layer thickness from images in panels **e** and **f** (left). Relative fluorescence intensity of expressed markers in panels **e** and **f** (right) (*n*=5). **h**, Production of extracellular fibrosis-associated proteins (IL-11, TGF-β, and COL1A1) in nFib and iFib cells cultured on Transwells or PDMS membranes for 5 days, measured by ELISA (*n*=5). **i**, Quantitative analysis of cellular orientation in nFib and iFib cultured on Transwells or PDMS membranes for 5 days. Left: % frequency of orientation angles. Right: % alignment profiles (*n*=5). Bars, 50 µm. \**p*<0.001, \*\**p*<0.01, \*\*\**p*<0.05.

The stiffness of the culture substrate significantly influences the morphological and functional characteristics of intestinal fibroblasts^35^. We evaluated the behavior of nFib and iFib cultured on substrates softer than conventional tissue culture flasks. When nFib were cultured for 5 days on a polyester nanoporous membrane in a Transwell (stiffness: 174.34±16.47 MPa; Supplementary Fig. 1a), they exhibited disorganized and weakly developed F-actin fibers, low collagen I (COL I) expression, and an absence of detectable α-SMA expression (Fig. 1e, upper). Addition of profibrotic molecules, such as interleukin 11 (IL-11; 10 ng ml^−1^)^36^ and transforming growth factor beta (TGF-β; 10 ng ml^−1^)^37^, to nFib cells cultured in Transwells resulted in substantial upregulation of COL I, but no detectable α-SMA expression (Supplementary Fig. 1e). In contrast, iFib displayed significantly elevated α-SMA expression co-localized with organized actin stress fibers (*p*<0.001) (Fig. 1e, lower; Supplementary Fig. 1d). On softer polydimethylsiloxane (PDMS) silicone membranes (stiffness: 20.67±1.13 kPa; Supplementary Fig. 1a), nFib cells showed marked reductions in F-actin expression (Fig. 1f, upper) and cell layer thickness (Fig. 1g, left), along with significantly decreased α-SMA (∼8.18-fold, *p*<0.001) and COL I expression (∼1.30-fold, *p*<0.01) compared to those on Transwell membranes (Fig. 1e, 1f). Notably, treatment with a profibrotic cocktail of IL-11 and TGF-β did not induce detectable α-SMA, although COL I expression was substantially increased (Supplementary Fig. 1e). Cross-sectional analysis revealed that iFib formed significantly thicker cell layers than nFib on both Transwell (∼1.50-fold; *p*<0.001) and PDMS substrates (∼1.46-fold; *p*<0.01) (Fig. 1g, left). Additionally, iFib secreted significantly higher levels of fibrosis-associated cytokines, including IL-11 and TGF-β, across both culture substrates, whereas the secretion of collagen type I alpha 1 (COL1A1) did not significantly differ between nFib and iFib (Fig. 1h). Morphologically, nFib exhibited random, non-aligned growth, whereas iFib consistently displayed pronounced directional alignment, with orientation angles skewed toward 90° regardless of substrate stiffness (Fig. 1i). These findings underscore the distinct biochemical and morphological characteristics of nFib and iFib, demonstrating that iFib robustly maintain disease-associated mechano-resistant features *in vitro*.

### Biomechanical cues amplify profibrotic signatures in inflammation-associated fibroblasts

We next investigated how mechanobiological cues, such as fluid shear stress and peristalsis-like cyclic mechanical strain, influence the cellular and molecular signatures of intestinal fibroblasts under conditions of epithelial barrier impairment. To recapitulate this pathophysiological microenvironment, we systematically decoupled and reconstituted these biomechanical stimuli into three distinct regimes: fluid shear stress alone (+Flow), cyclic mechanical strain alone (+Str), or a combination of both (+Flow, +Str), directly applied to fibroblasts cultured in a modular gut-on-a-chip system (Fig. 2a, Supplementary Fig. 2a). iFib cells were seeded in the upper microchannel of the gut-on-a-chip and subjected to controlled microfluidic flow (e.g., apical (AP), basolateral (BL), or both; 20 µl h^−1^ volumetric flow rate) and cyclic stretch induced by repetitive pneumatic vacuum suction (10% cell strain, 0.15 Hz frequency; Supplementary Fig. 2b) throughout the culture period (Fig. 2b).

**Fig. 2.**
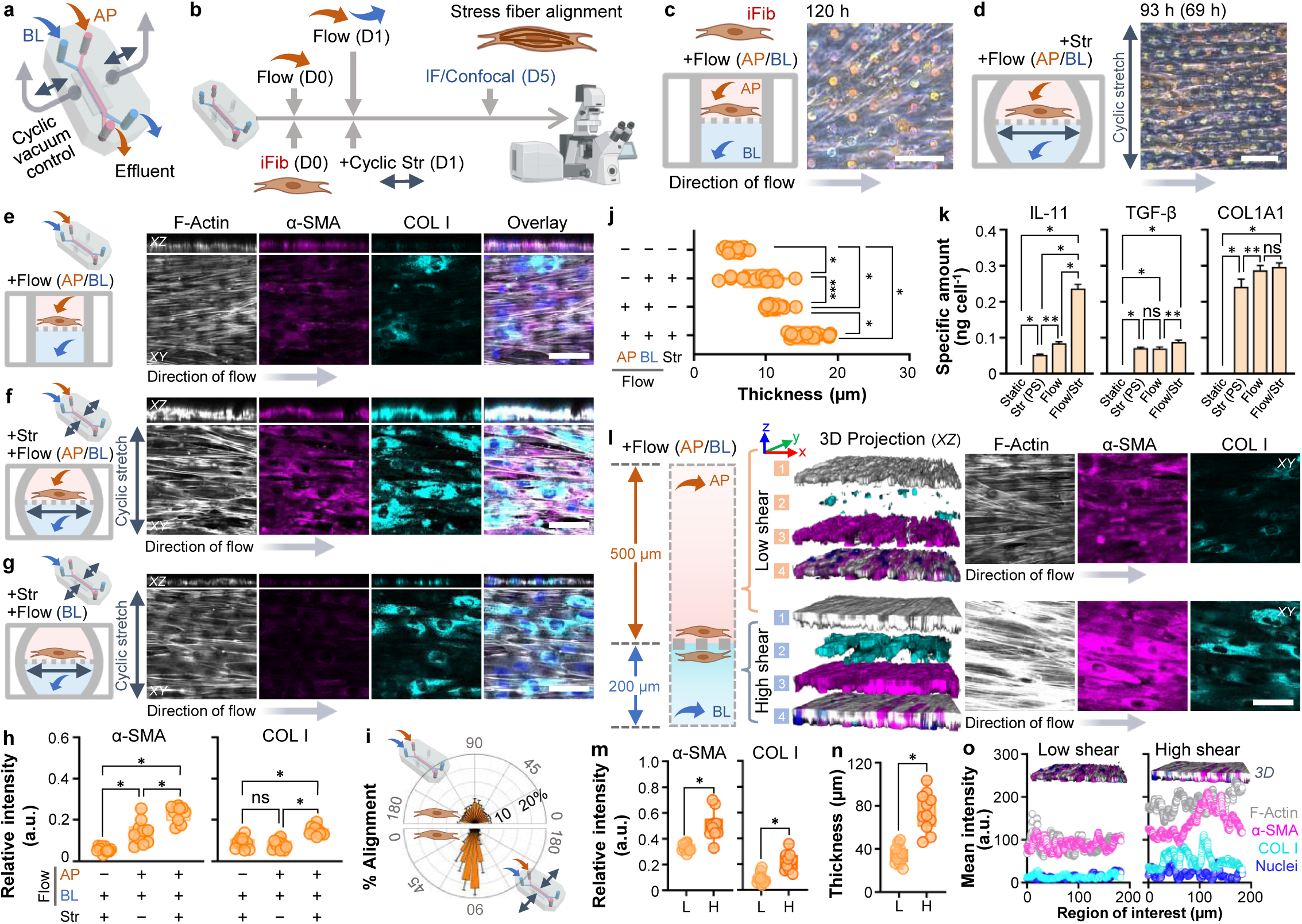
Fluid shear stress and cyclic mechanical strain promote profibrotic responses in inflammation-associated fibroblasts in a gut-on-a-chip. **a**, Schematic representation of the gut-on-a-chip system, illustrating the independent manipulation of biomechanical cues, including fluid shear stress applied to apical (AP; orange arrow) and basolateral (BL; blue arrow) and mechanical strain (double arrows). Grey arrows indicate the repetitive pneumatic suction to generate cyclic physical motions. **b**, Experimental workflow detailing the impact of biomechanical cues on iFib cells followed by immunofluorescence (IF) microscopy for phenotypic analysis such as the alignment of actin stress fibers with α-SMA. Experimental time points are designated as Day 0 (D0), Day 1 (D1), and Day 5 (D5). **c**, Phase-contrast micrograph illustrating robust growth of iFib under fluid shear stress (20 µl h^−1^ for 125 h) applied simultaneously to both apical (AP) and basolateral (BL) microchannels. **d**, A phase-contrast image showing perpendicular alignment of iFib cells subjected to cyclic mechanical strain (+Str) for 69 h under microfluidic dual flow (20 µl h^−1^; +Flow AP/BL; total culture time of 93 h). **e**, IF confocal images of iFib under microfluidic flow alone applied to both AP and BL microchannels (20 µl h^−1^; total culture time of 125 h), highlighting significant upregulation of profibrotic markers α-SMA (magenta) and COL I (cyan) compared to static controls on PDMS membranes shown in Fig. 1f. **f**, Combined microfluidic flow (20 µl h^−1^) and cyclic mechanical deformation (duration: 101 h) resulted in enhanced α-SMA and COL I expression, co-localization with actin stress fibers (F-actin; grey), cell elongation, and increased cell height. **g**, Microfluidic flow applied to the BL microchannel alone (20 µl h^−1^; total culture time of 125 h) under cyclic stretching caused iFib to align perpendicular to the stretch direction. **h**, Quantitative fluorescence intensity analysis of α-SMA and COL I expression across conditions shown in panels (**e**-**f**) (*n*=10). **i**, Orientation index analysis of iFib under cyclic deformation reveals significant cellular alignment with increased probability of orientation (*n*=10). **j**, Thickness of cell layers quantified from IF images in panels (**e**-**g**); static control data adapted from Fig. 1f (*n*=10). **k**, Production of extracellular fibrosis-associated proteins including IL-11, TGF-β, and COL1A1 in iFib cells cultured in a gut-on-a-chip (Str (PS), Flow, and Flow/Str) as well as static PDMS membrane (Static) for 5 days, measured by ELISA. PS, pseudo-static (*n*=3). **l**, Effect of fluid shear stress on iFib cultured on both sides of a PDMS porous membrane in the gut-on-a-chip, with low shear (∼0.00133 dyne cm^−2^) and high shear (∼0.00833 dyne cm^−2^) regimes induced by a flow rate of 20 µl h^−1^ for 125 h. **m**, Relative fluorescence intensity of α-SMA and COL I in iFib challenged to low (L) and high shear stress (H) in a gut-on-a-chip (*n*=3). **n**, Quantification of cell layer thickness under low (L) and high shear stress (H) regime from panel **l** (*n*=10). **o**, Mean fluorescence intensity of iFib cultured at low (L) and high shear stress (H) in a gut-on-a-chip as a function of detection location. XY, top-down views; XZ, vertical cross-sectional views. Bars, 50 µm. \**p*<0.001, \*\**p*<0.01, \*\*\**p*<0.05.

Under fluidic flow applied to both the AP and BL microchannels without mechanical stretching (“Fluidic”), iFib cells exhibited spindle-shaped morphologies with random orientations (Fig. 2c). However, when both fluid shear stress and cyclic stretch were applied simultaneously (“Dynamic”), the cells aligned perpendicularly to the direction of repetitive stretch even after 69 h of exposure (Fig. 2d). The “Fluidic” condition induced significantly elevated α-SMA and COL I expression (Fig. 2e, 2h) compared to static cultures (Fig. 1f, 1g). Under “Dynamic” conditions, iFib displayed a pronounced hypertrophic response characterized by increased cell layer thickness (Fig. 2f, 2j), upregulated α-SMA and COL I expression (Fig. 2f, 2h), and co-localization of actin stress fibers with α-SMA. Additionally, iFib cells demonstrated a strong orientational alignment in response to combined stimulation with fluid flow and cyclic mechanical strain (Fig. 2i). Conversely, under pseudo-static BL flow conditions without apical fluids shear, repetitive stretching alone did not significantly increase α-SMA expression (Fig. 2g, 2h), and cell layer thickness was markedly reduced (Fig. 2g, 2j), suggesting that apical fluid shear stress is a critical determinant for α-SMA induction in iFib. Mechanical strain, however, consistently enhanced COL I deposition across all conditions (Fig. 2f-2h), supporting its role in promoting ECM production. The secretion of profibrotic molecules, such as IL-11, TGF-β, and COL1A1, was maximized when fluid shear stress and mechanical strain coexisted (Fig. 2k). These results underscore the synergistic effects of biomechanical cues in amplifying profibrotic behaviors of iFib.

To assess the effect of fluid shear stress, iFib cells were seeded on a porous membrane in opposing orientations within a gut-on-a-chip, where the upper and lower microchannels had distinct heights of 500 and 200 µm, respectively. This height differential generated surface shear stress at approximately 1.33×^−3^ and 8.33×^−3^ dyne cm^−2^ in the upper and lower microchannels, respectively, resulting in a 6.25-fold shear stress gradient when a volumetric flow rate of 20 µl h^−1^ was applied to both channels. Under these conditions, iFib cells predominantly aligned in the direction of flow, exhibiting significantly increased α-SMA (∼1.53-fold; *p*<0.001) and COL I (∼2.46-fold; *p*<0.001) expression in the high-shear lower microchannel (Fig. 2l, 2m). Moreover, cell layer thickness was markedly greater (∼2.13-fold; *p*<0.001) in the high-shear regime (Fig. 2n), accompanied by enhanced actin stress fiber formation co-localized with α-SMA (Fig. 2o). These findings confirmed that iFib cells are resistant to biomechanical cues, exhibited a shear stress-dependent profibrotic phenotype, where elevated biomechanical stimuli drive enhanced ECM production and cytoskeletal remodeling *in vitro*.

### Apical shear stress drives atrophic damage in normal intestinal fibroblasts

Next, we examined how aberrant biomechanical stimulation influences the phenotypic and molecular perturbations on nFib using a gut-on-a-chip model. Exposure to apical shear stress (20 µl h^−1^) progressively induced morphological damage in nFib, resulting in significant cell detachment and apoptotic cell death over time (Fig. 3a, +Flow AP/BL), whereas cells remained viable under static conditions on the same PDMS membrane (Fig. 3a, Static). Notably, apical shear stress alone (+Flow AP) was sufficient to trigger cell detachment and apoptosis within 24 h (Fig. 3b), while basolateral shear stress (+Flow BL; 20 µl h^−1^) caused no comparable damage at 48 h and remained non-detrimental up to 120 h (Supplementary Fig. 3a). Under these conditions, BL flow did not induce upregulation of α-SMA or COL I expression (Fig. 3c, +Flow BL), even when combined with cyclic mechanical strain (Fig. 3c, +Flow BL, +Str). When cyclic stretch was applied with BL flow, nFib remained viable and exhibited a significantly increased cell layer thickness (∼1.68-fold compared to pseudo-static conditions; *p*<0.001). However, nFib cells did not display orientational alignment in response to cyclic stretch, with no difference compared to BL flow alone (Supplementary Fig. 3b). However, when fluid shear stress was applied to both channels (+Flow AP/BL) in conjunction with cyclic stretch, cell viability was maintained only up to 32 h, after which significant damage occurred, with over 70% of nFib cells exhibiting cell death (Fig. 3d). Under static conditions, the proliferative nFib population was 5.54±0.18%. However, even brief exposure (<18 h) to fluid shear stress significantly reduced proliferation (*p*<0.001), regardless of shear direction (1.90±0.15% for AP, 2.31±0.14% for BL, and 1.60±0.33% for both; Fig. 3e). While cyclic strain increased proliferation to 5.55±0.42% (*p*<0.001; Fig. 3e, +Flow AP/BL +Str), it was insufficient to maintain cell viability under fluid shear stress (Fig. 3d). These findings confirmed that direct exposure to apical shear stress is a crucial determinant of nFib cell fate, causing detachment and atrophic damage.

**Fig. 3.**
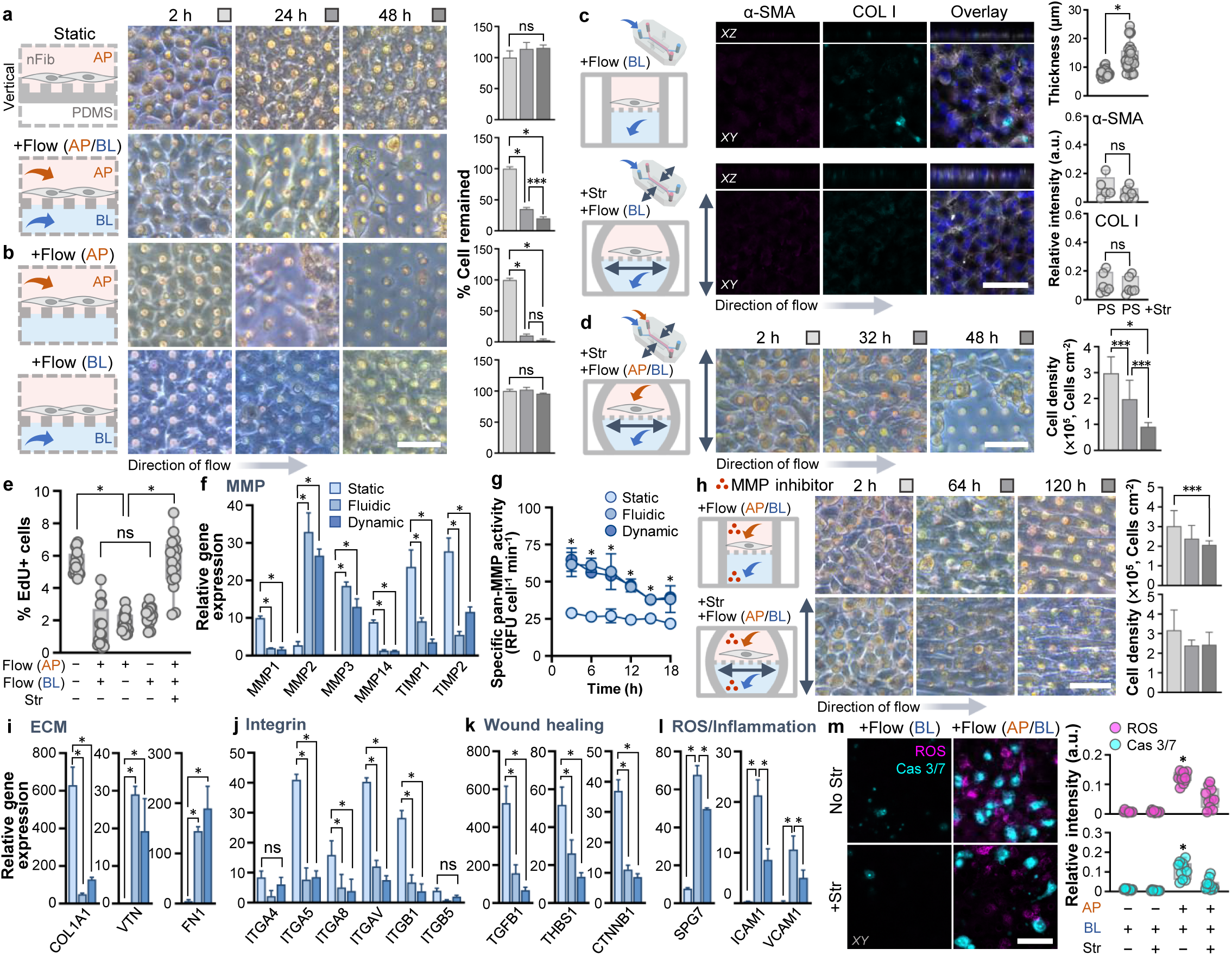
Normal intestinal fibroblasts undergo apoptotic injury under apical fluid shear stress in an MMP-dependent manner. **a,** nFib cells remained viable at 48 h when cultured on a PDMS membrane under static conditions (Static). In contrast, nFib cells cultured in a gut-on-a-chip subjected to fluid flow (20 µl h^−1^) on both AP and BL microchannels (+Flow, AP/BL) exhibited irreversible cell death within 48 h. Phase-contrast images were acquired at 2, 24, and 48 h. **b**, nFib cells were hypersensitive to apical fluid shear stress (+Flow, AP) but were resistant to basolateral shear stress (+Flow, BL). Quantification of remaining nFib cells, normalized to initial cell counts at 0 h, is shown in the right panel in **a** and **b** (*n*=5). **c**, Immunofluorescence micrographs at 94 h demonstrated that basolateral fluid shear stress did not induce cell death or expression of α-SMA (magenta) or COL I (cyan), irrespective of repetitive mechanical stretching (+Str). Overlaid images include signals of F-actin (gray) and nuclei (blue). The thickness of the cell layer (top right) and expression levels of α-SMA and COL I (bottom right) are quantified (*n*=3). **d**, Dynamic condition (+Flow, AP/BL & +Str) that involves constant fluid shear stress caused extensive nFib cell death, independent of cyclic stretching. Phase contrast images were taken at 2, 32, and 48 h, and cell numbers were quantified by counting nuclei (right panel) (*n*=3). **e**, Proliferative populations of nFib cells subjected to various flow conditions (AP vs. BL) and cyclic deformations (Str) were evaluated using EdU incorporation assays (*n*=4). **f**, Transcriptomic analysis revealed changes in matrix metalloproteinase (MMP)-associated genes (*MMP1*, *MMP2*, *MMP3*, *MMP14*, *TIMP1*, and *TIMP2*) under static, fluidic (dual flow, AP/BL), and dynamic (dual flow + cyclic stretching) conditions after 18 h, analyzed via qPCR (*n*=3). **g**, Pan-MMP catalytic activity was measured under static, fluidic, and dynamic conditions using a pan-MMP substrate^71^ (*n*=3). **h**, Treatment with a pan-MMP inhibitor (GM6001, 25 µM; indicated by red circles in the schematic) rescued nFib cell viability under both fluidic and dynamic conditions. Phase contrast images were acquired at 2, 64, and 120 h, with cell counts quantified by nuclei staining (*n*=3). **i-l.** Gene expression profiles of (**i**) extracellular matrix components (*COL1A1*, *VTN*, *FN1*), (**j**) integrins (*ITGA4*, *ITGA5*, *ITGA8*, *ITGAV*, *ITGB1*, *ITGB5*), (**k**) wound healing-related genes (*TGFB1*, *THBS1*, *CTNNB1*), and (**l**) the genes associated with oxidative stress and inflammation (*SPG7*, *ICAM1*, *VCAM1*) were examined (*n*=3). **m.** Cytoplasmic ROS (magenta) and apoptotic cell death (cyan) were visualized using a fluorogenic ROS detection reagent and a substrate for measuring caspase-3/7 activity. These phenomena were analyzed under apical and/or basolateral fluid shear stress (+Flow, BL vs. AP/BL) with or without cyclic stretching (+Str). Quantifications are presented in the right panels (*n*=3). Double arrows indicate the direction of cyclic stretch in panels of **c**, **d**, and **h**, while grey arrows represent medium flow direction in panels **b**-**d**, and **h**. Bars, 50 µm. \**p*<0.001, \*\**p*<0.01, \*\*\**p*<0.05.

Apical shear stress induced nFib cell shrinkage and detachment from the basement membrane, suggesting perturbation of genes involved in ECM regulation, cell adhesion, and stability. qPCR analyses revealed significant upregulation of matrix metalloproteinases (*MMP*) 2 and *MMP3* and downregulation of *MMP1*, *MMP14*, tissue inhibitors of metalloproteinases (*TIMP*) 1, and *TIMP2* under both fluidic (+Flow AP/BL) and dynamic (+Flow AP/BL +Str) conditions compared to static controls (Fig. 3f). The specific MMP activity peaked at 3 h in both Fluidic or Dynamic conditions, then progressively declined over time, whereas activity remained consistently low in static cultures regardless of the culture period (*p*<0.001; Fig. 3g). This elevated MMP activity, linked to excessive ECM degradation, likely contributed to atrophic cell death^38^. When treated with a pan-MMP inhibitor (GM6001; 25 µM), nFib cells were viable under shear stress conditions (+Flow AP/BL) for up to 120 h regardless of cyclic stretching, confirming the critical role of MMP regulation in maintaining nFib cell viability (Fig. 3h, Supplementary Fig. 3c). Notably, treatment with a pan-MMP inhibitor resulted in a pronounced orientational alignment under combined exposure to fluid flow and cyclic mechanical deformations as a function of time, compared to fluid flow alone (Supplementary Fig. 3d). Additionally, both Fluidic and Dynamic conditions led to heterogeneous ECM gene expression, with decreased collagen type I alpha 1 (*COL1A1*) and increased vitronectin (*VTN*) and fibronectin (*FN1*) levels (Fig. 3i). Integrin-associated genes, such as integrin subunit alpha 4 (*ITGA4*), *ITGA5*, *ITGA8*, *ITGAV*, integrin beta 1 (*ITGB1*), and *ITGB5*, were significantly downregulated under both Fluidic and Dynamic conditions (*p*<0.001; Fig. 3j). Genes related to homeostatic wound healing processes, including transforming growth factor beta 1 (*TGFB1*), thrombospondin-1 (*THBS1*), or catenin beta 1 (*CTNNB1*), were also markedly reduced in both Fluidic and Dynamic cultures compared to Static controls (Fig. 3k). In contrast, genes associated with oxidative stress (*SPG7*) and inflammatory adhesion responses, such as intercellular adhesion molecule 1 (*ICAM1*) and vascular cell adhesion molecule 1 (*VCAM1*), were significantly elevated under both Fluidic and Dynamic conditions (Fig. 3l). Exposure of nFib cells to apical fluid shear stress for 18 h markedly increased cytoplasmic reactive oxygen species (ROS) and caspase 3/7 (Cas 3/7)-mediated apoptotic activity (*p*<0.001) compared to pseudo-static controls (Fig. 3m, +Flow BL; Supplementary Fig. 3e & 3f). Notably, the application of cyclic mechanical strain significantly attenuated both ROS generation and Cas 3/7 activity (*p*<0.001), indicating a protective effect against shear-induced apoptosis. Together, these findings highlight fluid shear stress as a dominant biomechanical regulator of nFib viability, with an inverse relationship between apoptosis and proliferation determining mechanoresponsive cell fate.

### Intact epithelial barrier protects against shear-induced fibroblast injury

The intestinal epithelial barrier normally shields underlying fibroblasts from direct exposure to luminal fluid shear stress. We hypothesized that aberrant shear exposure could have detrimental effects on nFib. To verify this hypothesis, we employed a gut-on-a-chip model to simulate either an intact mucosal barrier (epithelial-fibroblast co-culture; Fig. 4a, schematic) or a compromised epithelial barrier (fibroblasts alone; Fig. 4b, schematic) under apical fluid flow. In the intact barrier model, normal intestinal organoid-derived epithelial cells were seeded into the apical microchannel and nFib cells into the basolateral microchannel on opposing sides of a porous membrane. Apical flow was applied at 100 µl h^−1^ for up to 72 h to challenge the system and assess epithelial protective effect against shear-induce fibroblast injury. Under intact barrier conditions, nFib labeled with CellTracker green maintained high confluency at 22 h (Fig. 4a) and stable vimentin expression at 72 h (Fig. 4c). In contrast, in the absence of epithelial coverage, nFib exhibited marked cell loss, weakened vimentin staining, prominent actin stress fibers, indicative of cytoskeletal reorganization and injury (Fig. 4b, 4c). Transepithelial electrical resistance (TEER) was significantly higher (*p*<0.001) in epithelial-fibroblast co-cultures compared to fibroblast monocultures (Fig. 4d), confirming the protective role of barrier integrity. Morphological analysis further revealed that nFib under intact epithelium displayed random F-actin organization (Fig. 4e, left) and stochastic alignment (Fig. 4f), whereas fibroblasts exposed directly to shear stress showed pronounced elongation and directional alignment (Fig. 4e, right). These findings elucidate the critical role of the epithelial barrier in mitigating the adverse effects of luminal shear stress on underlying fibroblasts and highlight its importance in maintaining intestinal tissue homeostasis.

**Fig. 4.**
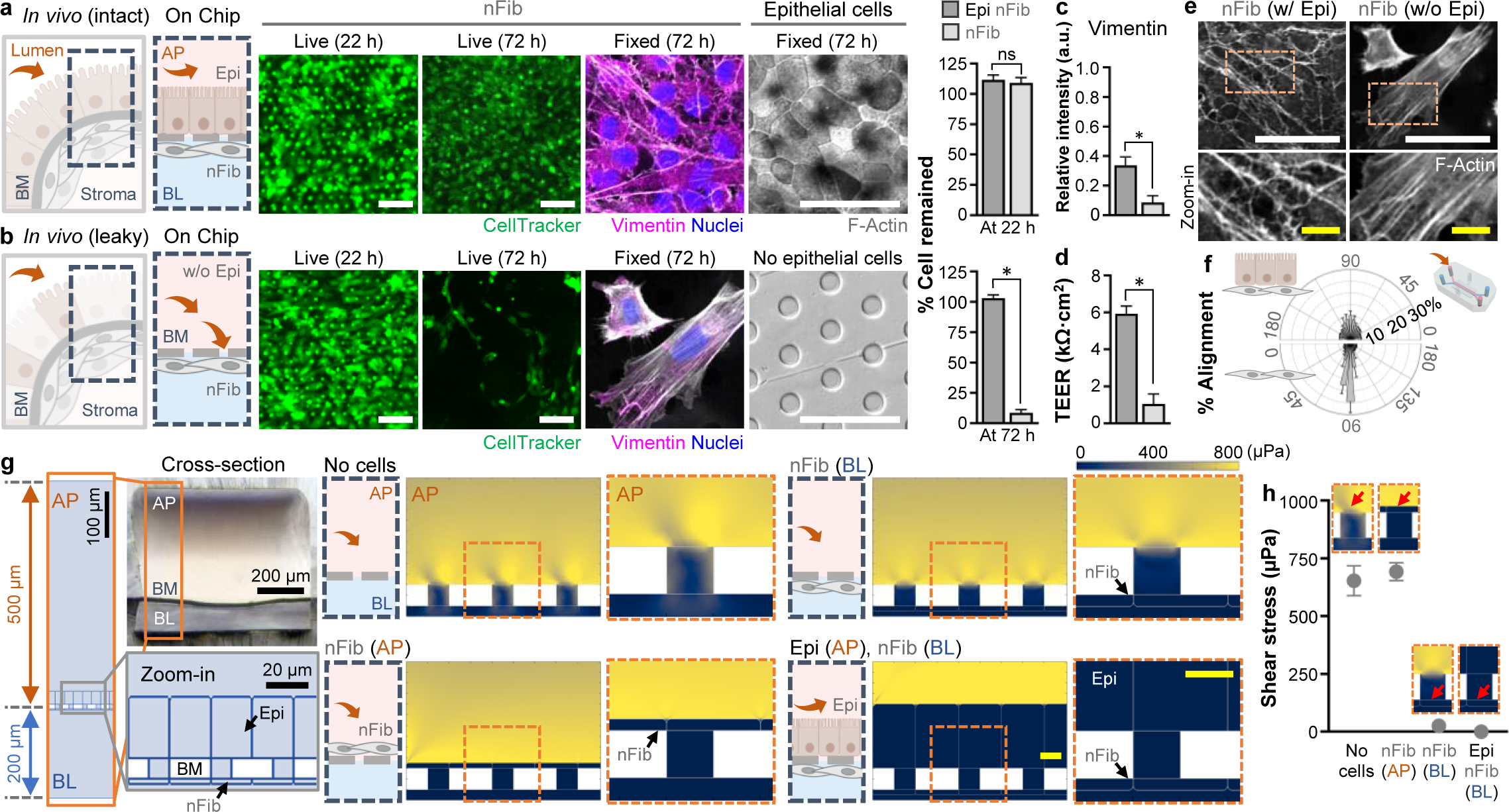
Intact intestinal epithelial barrier protects normal fibroblasts from fluid shear-induced injury in a gut-on-a-chip. **a,** Schematic representation of an intact epithelial-fibroblast interface *in vivo* (left) and its *in vitro* modeling using a gut-on-a-chip (right). Human intestinal epithelial cells from biopsy-derived organoids were co-cultured with nFib on opposite sides of a porous basement membrane (BM) to establish compartmentalization. Fluid flow at 100 µl h^−1^ was applied apically for up to 72 h. Live nFib morphology was tracked using green CellTracker dye, while fixed samples were characterized at 72 h via immunofluorescence confocal microscopy for vimentin (magenta), F-actin (gray), and nuclei (blue). nFib cell numbers were quantified by nuclear counting (*n*=3). **b**, Schematic of an impaired epithelial barrier *in vivo* (left) and its *in vitro* simulation on a gut-on-a-chip (right), where nFib were cultured alone on the lower side of the porous membrane. Live nFib morphology was monitored as in panel **a**, and remaining adherent nFib after 72 h of fluid flow were characterized by vimentin, F-actin, and nuclei immunostaining (*n*=3). **c**, Expression level of vimentin was quantified by comparing relative intensity shown in panel **a** and **b** (*n*=9). **d**, TEER measurements comparing barrier function between epithelial-fibroblast co-cultures and fibroblast mono-cultures (*n*=9). **e**, Comparison of actin cytoskeleton organization in nFib cultured in the presence of an epithelial layer (left) versus in mono-culture (right). White dashed boxes in the upper panels indicate regions magnified in the lower panels. **f**, Quantification of nFib cell alignment (orientation index) in the presence (upper) and absence (lower) of epithelial cells, as analyzed from the images shown in panel **d** (*n*=8). **g**, Schematic illustrations of a vertical section of the gut-on-a-chip microchannels (orange boxes), highlighting epithelial-nFib interface around the BM (gray boxes). Culture configurations include: No cells, nFib seeded on the AP side (nFib AP), nFib seeded on the BL side (nFib BL), and epithelial cells on the AP side with nFib on the BL side (Epi AP, nFib BL). Corresponding 2D simulations visually represent fluid shear stress distributions under each configuration. **h**, Quantification of the fluid shear stress experienced by nFib at their surface under the different configurations, indicated by red arrows in insets (No cells, *n*=129; nFib AP, *n*=2,785; nFib BL, *n*=279; Epi AP nFib BL, *n*=1,038). Bars: 50 µm (white); 10 µm (yellow). \**p*<0.001, \*\**p*<0.01, \*\*\**p*<0.05.

To characterize the influence of fluid dynamics on fibroblast responses, computational simulations were conducted to estimate fluid shear stress profiles at the cell surface, reflecting the geometry of the gut-on-a-chip (Fig. 4a, schematic; Supplementary Fig. 4a). Four configurations were modeled: a cell-free control (No cells), fibroblast monocultures positioned on the apical side (nFib AP) or basolateral side (nFib BL), and an epithelial-fibroblast co-culture (Epi AP, nFib BL). Simulations revealed the highest shear stress levels (692.06±0.15 µPa) on nFib cells directly exposed to luminal flow at the apical surface (Fig. 4g, 4h). In contrast, shear stress was negligible when nFib cells were positioned beneath the basement membrane, aligning well with linear flow rate profiles (Supplementary Fig. 4b). Indeed, apical fluid transmitted through 10 µm pores exerted substantially lower shear stress (25.76±0.80 µPa) on underlying fibroblasts. However, localized fluidic eddies, identified by surface velocity fields at steady state (Supplementary Fig. 4c, arrows), contributed to progressive nFib cell loss on the BL side. These findings confirmed that an intact epithelial barrier is both necessary and sufficient to preserve nFib viability under luminal shear conditions.

### Longitudinal exposure to luminal fluid shear induces the transition of normal fibroblasts into myofibroblast-like cells

As noted, when nFib cells were subjected to continuous fluid shear in a gut-on-a-chip for several days (Fig. 5a), the majority of nFib cells underwent apoptotic cell damage, leading to a substantial cell loss by ∼24 h. Interestingly, a few surviving populations among dying nFib cells formed sporadic cell aggregates, then progressively increased their size up to several hundred micrometers under the exposure to continuous fluid shear stress (Fig. 5b). Some aggregates began sprouting cells that spread across the surface of a porous membrane, demonstrating remarkable robustness under fluid shear stress (Fig. 5b, Zoom-in). Scanning electron microscopy (SEM) revealed that the surface of these aggregates was covered with massive collagen-like fibrils with multiple superposed cells underneath the fibrils (Fig. 5c, Supplementary Fig. 5a). The aggregates harvested at days 4 (D4) and 10 (D10) showed distinct phenotypic changes, where immunofluorescence confocal microscopy revealed that D4 aggregates lacked detectable expression of α-SMA and COL I across all vertical locations (Fig. 5d). In contrast, D10 aggregates exhibited strong α-SMA expression and mild COL I expression throughout their vertical cross-sectional structure (Fig. 5e). Notably, the sprouting populations on the membrane surface displayed the most remarkable α-SMA expression, overlapping with actin stress fibers, indicating a transition into mechanoadaptive myofibroblast-like cells (Fig. 5f). Quantitative analysis confirmed the progressive growth of these aggregates, with the average height of D10 aggregates (46.69 ± 2.07 µm) significantly greater (*p*<0.001) than that of D4 aggregates (28.36 ± 2.57 µm) (Fig. 5g). Fluorescence intensity profiles further validated the significant increase in both α-SMA and COL I expression from D4 to D10 (Fig. 5h, Supplementary Fig. 5b). Flow cytometry analysis also confirmed dramatic phenotypic shifts of nFib cells from the original state (nFib cells cultured in T75 flasks; Fig. 5i, T75) to the transient state (nFib cells challenged to microfluidic conditions in a gut-on-a-chip; Fig. 5i, D4 & D10) or from the reference state (iFib cells cultured in T75 flasks; Fig. 5i, iFib T75). We observed significant increases in fibrosis-associated markers, including α-SMA, fibronectin, and CD90 (Glycosylphosphatidylinositol-anchored glycoprotein, also known as Thy-1^39^) (Fig. 5i). Flow cytometry plots (Fig. 5i, left) and their quantification (Fig. 5i, right) showed a marked transition of nFib cells from their native state (T75) to myofibroblast-like aggregates at D10, reminiscent of iFib phenotypes. Notably, D10 aggregates exhibited significant elevation in α-SMA (∼4.72 folds) and fibronectin expression (∼3.02 folds) compared to nFib cells in T75 (*p*<0.001) but were not significantly different from iFib cells. Notably, CD90 expression was consistent across all cell types, reflecting its role as a general fibroblast marker. These findings collectively highlight the impact of fluid shear stress in driving the phenotypic reprogramming of normal fibroblasts into mechanoadaptive myofibroblast-like cells, providing insights into the mechanobiological processes underlying fibrosis-associated cellular dynamics.

**Fig. 5.**
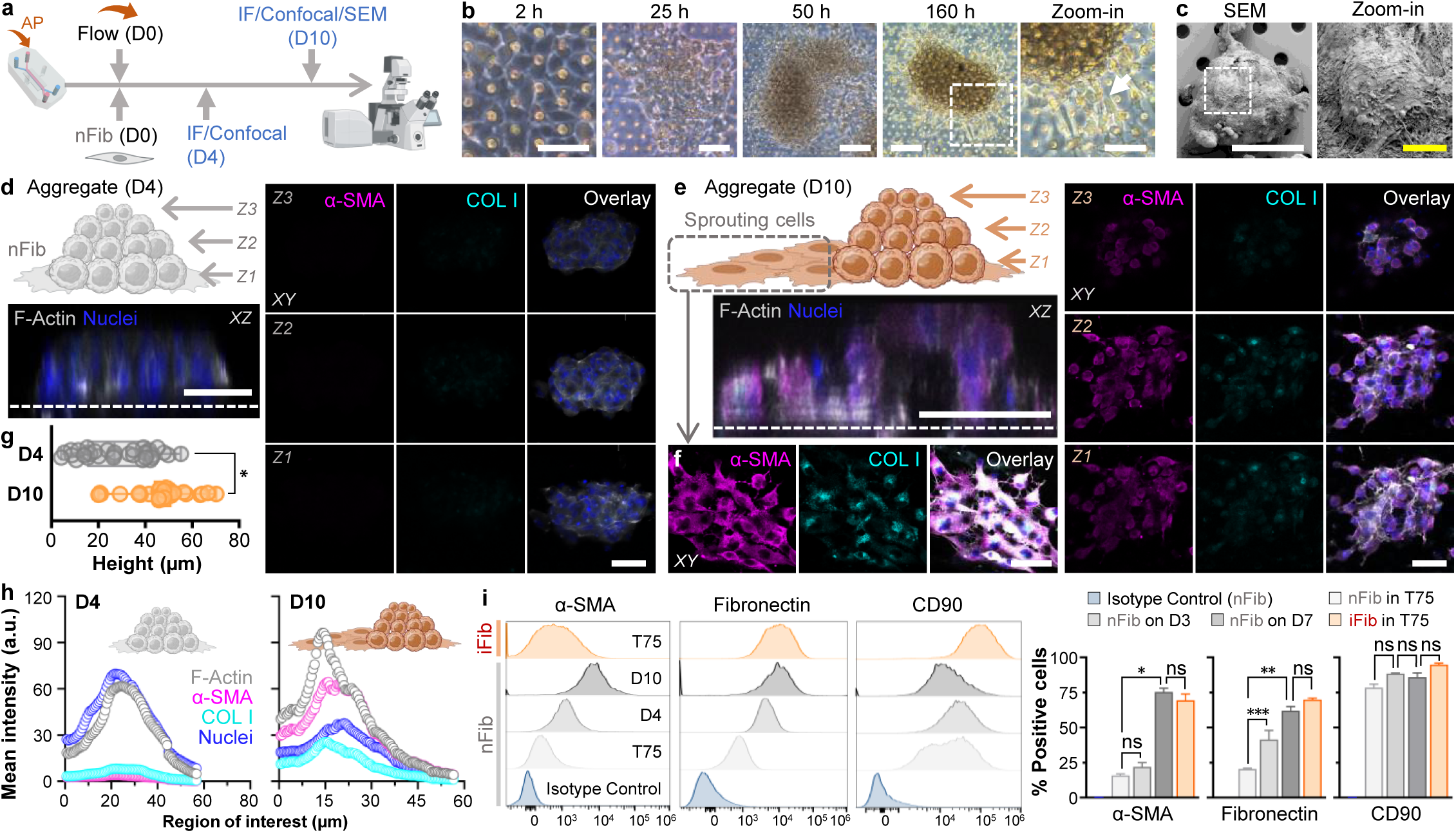
Longitudinal exposure to fluid shear induces the transition of normal fibroblasts to myofibroblast-like cells. **a**, Experimental workflow demonstrating the introduction of apical flow to nFib cells, followed by immunofluorescence (IF) confocal microscopy and scanning electron microscopy (SEM). Experimental time points are day 0 (D0), day 4 (D4), and day 10 (D10). **b**, Morphological changes in nFib cells exposed to apical shear stress (20 µl h^−1^) in a gut-on-a-chip. Phase contrast micrographs were taken at 2, 25, 50, and 160 h. White dashed square indicates the region of the Zoom-in view. The white arrow points to sprouting cells on the PDMS membrane. **c**, SEM images highlighting the formation of hemi-spherical nFib aggregates (left), with the Zoom-in view of the white dashed square (right). **d**, IF microscopy of nFib aggregates on D4, showing poor expression of α-SMA (magenta) and COL I (cyan). Overlaid images depict F-actin (grey) and nuclei (blue). Z1, Z2, and Z3 represent the Z-position of the corresponding XY views. **e**, IF images of fibrotic markers α-SMA and COL I in nFib aggregates on D10, with corresponding XY views at Z1, Z2, and Z3 positions. Overlaid images show F-actin (grey) and nuclei (blue). **f**, IF views of α-SMA and COL I in sprouting nFib cells on the PDMS surface from aggregates on D10. **g**, Quantification of nFib aggregate height between D4 and D10 (*n*=33). **h**, Mean intensity profiles of α-SMA, COL I, F-actin, and nuclei in aggregates at Days 4 and 10. **i**, Flow cytometry histograms comparing the profile of nFib cells from cultures in T75 flasks, aggregates at D4 and D10, and iFib cultured in T75 flasks, focusing on fibrosis-associated markers such as α-SMA, fibronectin, and CD90. Quantification is shown in the right panel (*n*=3). XY: top-down views; XZ: vertical cross-sectional views. Bars: 50 µm (white), 10 µm (yellow).

### Mechanoadaptive fibroblasts from shear-induced aggregates exhibit profibrotic traits

To investigate the cellular and molecular characteristics of the fibroblasts within shear-induced aggregates, D10 aggregates were enzymatically dissociated using trypsin, and the resulting cells were expanded through serial passages in T75 flasks (Fig. 6a). The isolated mechanoadaptive fibroblasts (nFib_MA_) displayed a spindle-shaped, elongated morphology reminiscent of myofibroblasts, with a marked anisotropic alignment. Quantitative image analysis revealed that nFib_MA_ exhibited a significantly higher orientation index compared to nFib (*p*<0.001, Supplementary Fig. 6a). Flow cytometry analysis confirmed that nFib_MA_ exhibited significantly elevated expression both α-SMA and fibronectin among CD90+ and vimentin+ cell populations, aligning closely with the expression profile of iFib (Fig. 6b, Supplementary Fig. 6b, 6c). Structural assessment via SEM revealed that both iFib and nFib_MA_ had roughened surfaces with multiple microvilli-like protrusions, a stark contrast to the smoother morphology of nFib (Fig. 6c). Additionally, atomic force microscopy (AFM) quantitatively confirmed that both surface roughness (Fig. 6d) and average cell height (Fig. 6e) were significantly elevated (*p*<0.001) in both iFib and nFib_MA_ compared to nFib, with no substantial distinction between iFib and nFib_MA_. Moreover, AFM-based stiffness measurements demonstrated that Young’s modulus values of iFib, nFib_MA_, D4, and D10 aggregates were significantly increased relative to nFib, indicating a progressive stiffening response to mechanical stress (Fig. 6f).

**Fig. 6.**
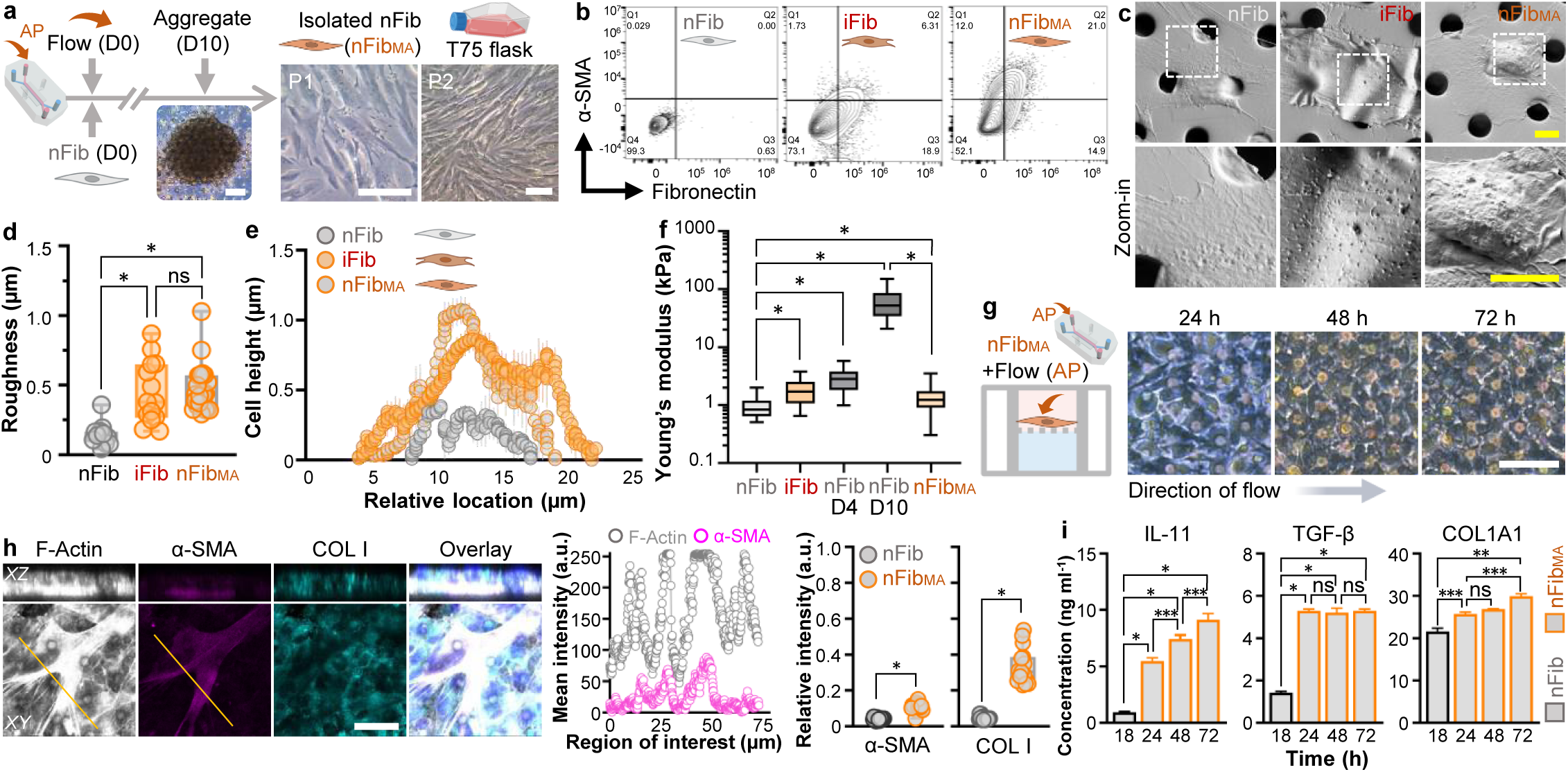
Mechanoadaptive fibroblasts isolated from shear-induced aggregates exhibit profibrotic phenotypes and behaviors. **a**, A workflow illustrating the formation of nFib aggregates under continuous apical flow (20 µl h^−1^) in a gut-on-a-chip, followed by isolation and expansion of mechanoadaptive fibroblasts (nFib_MA_) in T75 flasks. Representative phase contrast micrographs show nFib_MA_ morphology at passages 1 (P1) and 2 (P2) when cells reached confluence. **b,** Flow cytometric comparison of nFib, iFib, and nFib_MA_ cultured on T75 flasks, highlighting the expression of profibrotic markers α-SMA and fibronectin. **c**, SEM views of nFib, iFib, and nFib_MA_ cells cultured on a PDMS membrane (top) and magnified views of the regions outlined by white dashed squares (bottom). **d**, Surface roughness profiles of nFib, iFib, and nFib_MA_ cells derived from SEM images in panel **c**, quantified using AFM (*n*=300). **e**, Height of nFib, iFib, and nFib_MA_ cells cultured statically on PDMS membranes (*n*=15). **f**, Young’s modulus measurements of nFib, iFib, nFib_MA_ cells, and nFib aggregates (D4 and D10) under static culture conditions (*n*=500). D4 and D10 aggregates were generated under microfluidic conditions (20 µl h^−1^). **g**, Phase contrast micrographs of nFib_MA_ cultured in a gut-on-a-chip under apical flow (20 µl h^−1^) for up to 72 h. **h**, IF microscopy reveals the expression of profibrotic α-SMA (magenta) and COL I (cyan) on nFib_MA_ cells cultured in a gut-on-a-chip under fluid shear stress for 95 h. An overlay image includes F-actin (gray) and nuclei (blue) signals. The mean fluorescence intensity of the line scan applied on F-actin and α-SMA images (orange lines) as a function of the region of interest (middle panel). Quantitative expression levels are presented on the right panel (*n*=3). **i**, Quantification of secretory fibrosis-associated markers, including IL-11, TGF-β, and COL1A1 from nFib and nFib_MA_ cells cultured in a gut-on-a-chip for 72 h (*n*=3). Bars: 50 µm (white), 10 µm (yellow). \**p*<0.001, \*\**p*<0.01, \*\*\**p*<0.05.

To further examine how nFib_MA_ behave under sustained fluid shear stress, we cultured them within a gut-on-a-chip microdevice and exposed them to continuous microfluidic flow at 20 µl h^−1^ for 72 h. Under these conditions, no significant cell detachment was observed (Fig. 6g). Immunofluorescence confocal microscopy demonstrated that nFib_MA_ exhibited pronounced α-SMA expression, which strongly co-localized with actin stress fibers, alongside a marked increase in COL I deposition relative to nFib (Fig. 6h). Further biochemical analyses revealed that profibrotic cytokines and extracellular matrix components, including IL-11, TGF-β, and COL1A1, were significantly upregulated in nFib_MA_ compared to nFib (Fig. 6i). Notably, IL-11 secretion exhibited a time-dependent increase, whereas TGF-β and COL1A1 levels remained relatively stable or showed a less pronounced change over time. This result suggests that IL-11 plays a central role in mediating the profibrotic response under prolonged exposure to fluid shear stress.

## Discussion

Intestinal fibrosis is a major pathological complication of IBD, yet its initiating mechanisms and regulatory pathways remain poorly understood. Emerging evidence suggests that mucosal damage serves as the primary site for the onset of fibrosis, implicating mucosa-associated cells, such as intestinal epithelial cells and fibroblasts, and microenvironmental factors, including mechanostimulatory forces, as critical modulators of early profibrotic processes^4, 40,41^. However, the absence of physiologically relevant human models has limited mechanistic investigations into how the initial fibrogenic microenvironment drives pathological progression.

In this study, we leveraged a microengineered human gut-on-a-chip platform to recreate the biomechanically dynamic intestinal microenvironment and elucidate its role in initiating early fibrotic events. We established experimental models featuring fibroblasts cultured with either an intact or disrupted epithelial barrier, simulating normal and impaired mucosal conditions, respectively. Unlike conventional static cultures, which fail to capture the dynamic and multifactorial nature of intestinal fibrosis, this bioengineered system integrates controlled luminal flow and peristalsis-like mechanical deformations, enabling precise modulation of mechanotransductive signaling. Given that peristalsis-driven mechanical forces are intrinsic to gut physiology, their omission in traditional culture systems represents a major limitation in accurately modeling fibrosis pathogenesis. These findings underscore the necessity of utilizing physiologically dynamic in vitro models, such as the gut-on-a-chip, to dissect the complex interplay between mechanical forces and fibrotic remodeling in a human-relevant context.

Notably, the modular design of this model allows for the independent interrogation of fluid shear stress and cyclic mechanical strain, facilitating the delineation of their respective contributions to fibroblast activation, which has yet to be challenging in animal models and human clinical studies. A key advantage of this approach is its ability to modulate biological complexity. *In vivo*, the intestinal microenvironment is highly complex, with SMCs^42^ contributing both mechanical forces and biochemical signals that regulate fibrosis^43^. Because SMCs not only generate contractile strain but also secrete fibrogenic factors, it is difficult in animal models to separate the effects of mechanical strain from molecular signaling. The gut-on-a-chip system overcomes this challenge by recreating peristalsis-like mechanical deformations without the presence of SMCs, allowing the specific effects of cyclic strain on fibroblast behavior to be studied in isolation. This physiologically relevant, yet reductionist, platform provides a controlled way to investigate how mechanical forces drive early fibrotic processes - insights that are difficult to obtain from conventional *in vivo* models.

When utilizing patient-derived fibroblasts, it is essential to ensure that their disease-intrinsic characteristics are stably maintained *in vitro*. Most previous studies on fibroblast behavior in fibrosis have focused on responses to ECM remodeling and tissue mechanoregulation, particularly matrix stiffness^44, 45^. In this context, we observed distinct morphological differences between nFib and iFib in response to varying substrate stiffness across different culture formats. AFM measurements of substrate Young’s modulus aligned well with previously reported values: T-flask (∼3 GPa^46^), polyester nanoporous membrane (<2 GPa^47^), and PDMS membrane (<1 MPa^48^). Notably, the substrates used in this study exhibit significantly higher stiffness compared to *in vitro* hydrogels (∼0.1 kPa^49^) and *in vivo* IBD tissue (∼1 kPa^50^), a key factor to consider when interpreting mechanobiological responses.

Consistent with prior observations, stiffer substrates promoted fibroblast elongation and alignment^51^, and induced α-SMA expression in iFib, a hallmark of myofibroblast differentiation. In contrast, nFib exhibited no detectable α-SMA expression under identical conditions. This distinct mechanophenotypic behavior was consistently maintained across all culture formats, underscoring the reliability of patient-derived fibroblasts for studying disease-relevant mechanobiological responses.

Dysregulated bowel motility is one of the characteristics of IBD, yet its influence on early fibrotic remodeling and the underlying cellular mechanisms remains insufficiently understood^19–21^. Our microfluidic and stretchable gut-on-a-chip replicates physiologically relevant mechanical cues, in contrast to static cultures, which lack bowel motility and are mechanically abnormal. We found that fluid shear stress promoted hypertrophy in iFib, increased fibroblast layer thickness, and enhances myofibroblast-like activation, as indicated by increased α-SMA, TGF-β, and IL-11. In contrast, nFib remained were highly sensitive to the same mechanical cues, exhibiting limited activation and instead undergoing significant cell death. These results demonstrate that iFib exhibit heightened mechanosensitivity, undergoing cytoskeletal reorganization and activating profibrotic signaling pathways in response to shear stress and cyclic mechanical strain. Notably, the fibrotic response was further amplified when cyclic stretch was combined with fluid flow, exacerbating myofibroblast-like activation. In comparison, nFib showed minimal response, suggesting that inflammation-associated fibroblasts with predisposition to fibrotic activation are more susceptible to dynamic mechanical forces, thereby amplifying early fibrotic progression. Together, these findings highlight the limitations of conventional static cultures and emphasize the need for physiologically dynamic models to accurately study the biomechanical drivers of intestinal fibrosis in IBD.

We experimentally demonstrated that nFib cells are highly susceptible to fluid shear stress, particularly when applied apically. This stimulation results in atrophic cell detachment followed by apoptotic cell death within approximately 24 h, even at very low shear stress (∼0.00133 dyne cm^−2^). This hyper-sensitivity to fluid shear stress was consistent across different culture substrates and ECM coating conditions. Histologically, intestinal fibroblasts are localized beneath the epithelial barrier, where in healthy intestinal mucosa, they experience only minimal interstitial fluid shear in the sub-epithelial area or lamina propria. Our experimental model, therefore, replicated a pathological condition in which a disrupted epithelial layer mimics disease-associated sites (e.g., mucosal damage in IBD), within a controlled microphysiological environment. *In vivo*, epithelial barrier impairment triggers homeostatic processes that accompany wound healing, including myofibroblast activation through mechanical cues (e.g., cytoskeleton reorganization into stress fibers^52^), biochemical signaling (e.g., TGF-β release, ECM deposition^53^), and functional differentiation (e.g., α-SMA expression^54^). These processes were successfully replicated *in vitro* using disease-associated iFib in the gut-on-a-chip model. In contrast, nFib cells exhibited extreme sensitivity to prolonged fluid shear stress, resulting in apoptotic cell death within 24 h. As such, we conducted short-term (up to 18 h) fluid shear stress exposure, followed by comprehensive morphological, transcriptomic, and cellular assessments.

Cyclic stretching is known to promote fibroblast proliferation through mechanosensitive pathways such as YAP/TAZ signaling^55^ or integrin-FAK signaling^56^, a finding consistent with our results (Fig. 3e). However, despite this proliferative response, mechanical deformation alone was insufficient to mitigate the shear-induced apoptotic damage in nFib cells. To investigate the underlying molecular mechanisms, we first examined the expression of MMP-associated genes and their catalytic activity. When nFib cells were subjected to apical fluid shear stress for 18 h, we observed a significant upregulation of *MMP2* and *MMP3*, alongside a downregulation of their antagonists, including *TIMP1* and *TIMP2*. This transcriptomic alteration resulted in a marked increase in pan-MMP catalytic activity under microfluidic conditions, whereas activity in static controls remained largely unchanged regardless of culture duration. We hypothesize that the upregulation of MMPs in response to shear stress leads to the degradation of ECM components, such as collagen and fibronectin, weakening the structural support for fibroblasts. This degradation reduces focal adhesion stability, which can result in detachment-induced apoptosis (anoikis) or cellular damage^57^. MMP-mediated ECM degradation disrupts the anchoring points necessary for efficient force transduction. Without a stable ECM, fibroblasts are unable to properly polarize their cytoskeleton in response to cyclic strain, preventing alignment^58, 59^. This hypothesis was further validated by the use of an MMP inhibitor (GM6001), which preserved ECM integrity by inhibiting excessive ECM degradation. With an intact matrix, fibroblasts maintained stable focal adhesions, enabling proper mechanotransduction. Under these conditions, fibroblasts effectively sensed and adapted to mechanical stimuli, aligning their actin filaments and microtubules perpendicular to the direction of cyclic stretching.

Under dynamic shear stress and strain, intestinal fibroblasts undergo extensive ECM remodeling, characterized by increased vitronectin (*VTN*) expression and reduced integrin levels (*ITGA5*, *ITGA8*, *ITGAV*, *ITGB1*), suggesting a decoupling of ECM production from cell adhesion. *VTN* upregulation can serve as a protective ECM-stabilizing response^60^, while integrin downregulation suppresses mechanotransduction, potentially shifting fibroblasts toward a quiescent or apoptotic state^61^. This balance could mitigate excessive ECM deposition and fibrosis by reducing TGF-β signaling and myofibroblast activation but may also impair fibroblast adhesion, migration, and mucosal healing, contributing to chronic tissue damage in IBD^62^. The downregulation of *COL1A1* suggests reduced matrix rigidity and fibroblast activation, potentially mediated by inhibition of TGF-β (*TGFB1*) and Wnt/β-catenin (*CTNNB1*) signaling. Meanwhile, fibronectin (*FN1*) upregulation under dynamic conditions may be a compensatory mechanism to restore ECM integrity and cell adhesion^63^. Thrombospondin-1 (*THBS1*), a key activator of latent TGF-β and driver of fibrosis^64^, is significantly downregulated under mechanical stimulation, reducing TGF-β activation and shifting fibroblasts toward a less fibrotic state. These findings highlight the anti-fibrotic potential of mechanobiological cues in regulating ECM homeostasis and suggest *THBS1* as a therapeutic target for intestinal fibrosis in IBD. Additionally, spastic paraplegia 7 (*SPG7*), a mitochondrial matrix protein involved in the regulation of mitochondrial permeability transition pore (mPTP), plays a key role in oxidative stress and apoptosis by facilitating prolonged mPTP opening, leading to cytochrome c release and subsequent caspase activation^65^. We found that mechanical stress under flow conditions has been shown to increase mitochondrial activity and ROS production in cells with elevated *SPG7* expression, correlating with heightened oxidative stress observed in fibrotic tissues. In our model, fluidic flow alone induced the highest upregulation of *SPG7*, whereas the addition of cyclic mechanical strain significantly suppressed its expression. This pattern of *SPG7* regulation corresponded with changes in inflammatory gene expression (*ICAM1*, *VCAM1*), cytoplasmic ROS levels, and caspase-3/7 (Cas3/7) activity, further emphasizing the critical role of mechanical microenvironments in modulating fibroblast oxidative stress responses and inflammatory activation.

Our findings confirm that an intact epithelial barrier is essential for protecting shear-sensitive nFib cells. When this barrier is compromised, nFib cells experience severe damage and undergo apoptosis. Interestingly, a subset of fibroblasts exhibited persistence under shear stress, forming atypical cellular aggregates. SEM analysis revealed that these aggregates consist of multiple fibroblasts embedded within a dense, fibrillar network, a previously unreported unique morphological feature. This dome-shaped aggregation and intricate fibril structure may create a protective niche, enabling fibroblasts to resist shear stress and transition into myofibroblast-like activated cells. This process likely reflects localized excessive ECM deposition, potentially representing an early stage of fibrosis in IBD. AFM analysis demonstrated that the stiffness of these aggregates on days 4 and 10 closely matched the tissue stiffness observed in IBD fibrosis patients^35^. Mechanoadaptive fibroblasts (nFib_MA_) isolated from these aggregates exhibited key myofibroblast-like characteristics, including increased hypertrophy, rough cell surfaces, elevated stiffness, and α-SMA expression aligned with actin stress fibers. Notably, nFib cells did not express detectable α-SMA under any conditions, including variations in culture substrate, mechanical stimulation, short-term fluid shear stress, or exposure to profibrotic factors such as IL-11 or TGF-β. Flow cytometry further confirmed that nFib_MA_ cells share markers associated with inflammation-associated fibroblasts, indicating that prolonged mechanodynamic stimulation induces a stable reprogramming of fibroblast phenotype. The morphological and molecular characteristics of nFib_MA_ remained unchanged even after the removal of mechanical stress, indicating irreversible mechanogenomic alterations in nFib through genomic reprogramming. Further studies are needed to determine whether these persistent transcriptomic and phenotypic changes are accompanied by genomic mutations, chromosomal alterations, or epigenetic modifications.

Despite the novel findings, our study has several limitations. First, we primarily utilized a limited number of fibroblast sources, necessitating validation with additional normal intestinal fibroblast populations to confirm the shear-induced transition. Second, our model included only two mucosal cell types, epithelial cells and fibroblasts. To better replicate the mucosal microenvironment and elucidate intercellular crosstalk in early fibrogenesis, future studies should incorporate additional cell types such as smooth muscle cells, enteric neurons, intestinal microvascular endothelial cells, and gut microbiota. Third, while this study highlights the role of biomechanical cues as inducers of fibrosis, their potential suppressive effects were not extensively explored. Further investigations should address this aspect to provide a more comprehensive understanding of fibrosis regulation. Finally, our gut-on-a-chip model employs a PDMS-based membrane, which differs in stiffness from the native basement membrane and surrounding ECM. Incorporating patient-derived biomaterials to create a physiologically relevant basement membrane and ECM could enhance the fidelity of disease modeling.

The gut-on-a-chip technology enabled the discovery of a novel mechanoadaptive reprogramming mechanism in normal intestinal fibroblasts, which are highly sensitive to fluid shear stress. This model allowed precise interrogation of intrinsic cellular properties and mechanostimulatory factors, both independently and in combination, within a controlled microenvironment. The establishment of a versatile platform that integrates mechanical cues, cellular interactions, and spatial organization facilitates a comprehensive understanding of fibrosis progression^66^. By leveraging microtechnology and patient-derived samples, this system is uniquely positioned to identify disease-specific manifestations and pinpoint the early triggers of profibrotic processes, paving the way for novel therapeutic targets. By accurately replicating the dynamic interplay between mechanical forces and cellular adaptation, this platform provides an invaluable tool for dissecting the complex etiology of intestinal fibrosis, insights that are particularly challenging to obtain from clinical studies, especially in patients with refractory fibrotic disease. Quantitative functional assays, combined with genomic, transcriptomic, and morphological analyses, will further support these findings and advance translational precision medicine.

## Methods

### Device fabrication

The gut-on-a-chip device was fabricated using standard photolithography and soft lithography techniques, as previously described^26, 28^. Polydimethylsiloxane (PDMS; Sylgard 184, Dow Corning) was prepared by mixing the base elastomer and curing agent at a 10:1 (w/w) ratio, then cast into SU-8-patterned silicon molds to form the upper and lower microchannel layers. After curing at 60 °C for 6 h, the PDMS layers were demolded to yield channels with dimensions of 1 mm width, 1 cm length, and heights of 500 μm (upper channel) and 200 μm (lower channel), respectively. A porous, elastic PDMS membrane (10 μm pore diameter, 25 μm center-to-center spacing, 10 μm thickness; prepared with a 10:1 base-to-curing-agent ratio) was fabricated following established protocols^26^ and placed between the two channel layers to mimic the intestinal basement membrane. Each microchannel was connected to programmable syringe pumps via silicone tubing (inner diameter: 1/32”, outer diameter: 3/32”; Tygon 3350, Beaverton) and 90°-bent blunt-end stainless steel needles (18G; Kimble Chase) for fluidic control. Two side chambers flanking the central cell culture channel were integrated into the device to enable the application of cyclic mechanical strain.

### Cell Cultures

Primary human intestinal fibroblasts were used to model healthy and inflammation-associated stromal responses. Normal intestinal fibroblasts (nFib; cat. no. 2920; ScienCell) and patient-derived inflammation-associated fibroblasts (iFib) isolated from the inflamed colonic tissue from a patient with ulcerative colitis were cultured in a fibroblast culture medium (Dulbecco’s Modified Eagle Medium, DMEM; Gibco) supplemented with 20% (v/v) heat-inactivated fetal bovine serum (FBS; Gibco), 1% (w/v) L-glutamine, and antibiotics (100 U ml^−1^ Penicillin and 100 µg ml^−1^ Streptomycin; Gibco). Cells (passage number <6) were maintained in 75 cm^2^ tissue culture flasks at 37 °C in a humidified 5% CO_2_ incubator and passaged upon reaching ∼90% confluency.

The iFib cells were isolated from surgically resected de-identified colonic specimens of a UC patient with severe inflammatory injury (S-241113-00981) under an approved protocol by the Institutional Review Board (IRB; 06-050) of Cleveland Clinic Tissue Center. Tissues were rinsed in phosphate-buffered saline (PBS, Ca^2+^ and Mg^2+^ free; Gibco), minced into fragments (<0.2 mm), and enzymatically digested in Hank’s Balanced Salt Solution (HBSS; Gibco) containing 10% (v/v) FBS, antibiotics cocktail (100 units ml^−1^ Penicillin, 100 μg ml^−1^ Streptomycin, and 0.25 μg ml^−1^ Amphotericin B; cat. no. 10-378-016; Gibco), and a cocktail of collagenases: type I (100 U ml^−1^; cat. no. 1639), type II (100 U ml^−1^; cat. no. 1764), and type IV (100 U ml^−1^; cat. no. C5138; all from Sigma-Aldrich). Digestion was carried out at 37 °C for up to 3 h in a shaking water bath. The resulting cell suspension was triturated gently, filtered through a 70 μm cell strainer (Celltreat Scientific Products), and centrifuged (300×*g* for 10 min, at 4 °C) to isolate fibroblasts. The purified iFib population was expanded and used for downstream experiments.

Normal human intestinal organoids were derived from deidentified biopsy samples obtained from a healthy donor (62-year-old Caucasian male) under an approved IRB (2017-06-0114) from Dell Medical School, The University of Texas at Austin. Informed consent was obtained from the donor prior to tissue collection. Biopsy specimens were acquired from the right colon during a routine endoscopic examination performed for clinical evaluation. Following tissue processing, intestinal crypts were isolated and embedded in Matrigel (Corning; cat. no. CB-40234) and maintained in a defined organoid culture medium as previously described^26^. Organoids were cultured at 37 °C in a humidified 5% CO_2_ incubator, with medium changes every other day. Passaging was performed every 7 days to maintain long-term culture and proliferation.

### Microfluidic cultures

Prior to cell seeding, the gut-on-a-chip devices were sterilized by sequentially introducing 70% (v/v) ethanol into the microchannels, followed by incubation at 60 °C for 6 h and ultraviolet-ozone treatment using a UVO Cleaner 342 (Jelight Company Inc.) for 40 min. Surface functionalization was performed to enhance ECM adhesion by incubating the microchannels with 1% (w/v) polyethyleneimine (Sigma-Aldrich, cat. no. 408700) for 10 min, followed by rinsing with sterile deionized water. Subsequently, 0.1% (v/v) glutaraldehyde (Electron Microscopy Sciences, cat. no. 16320) was introduced for 30 min at room temperature, after which the channels were thoroughly flushed with deionized water, then completely dried. To prepare the ECM coating, an ice-cold solution containing collagen I (0.03 mg ml^−1^; Gibco, cat. no. A10483-01) and Matrigel (0.3 mg ml^−1^; Corning) was injected into the channels and incubated at 37 °C for 1 h to promote matrix adsorption. Prior to fibroblast seeding, the upper microchannel was perfused with fibroblast culture medium at 20 μL h^−1^ for 12 h to precondition the matrix-coated surface. For static control experiments, a flat PDMS membrane was placed in a glass-bottom dish (Matsunami Glass Ind.; cat. no. D1113OH), sterilized with 70% (v/v) ethanol, dried at 60 °C for 6 h, and treated with UVO for 40 min, then followed the identical protocol for surface functionalization. For Transwell-based static cultures, nanoporous polyester membrane inserts (pore size: 0.4 μm; Corning) were subjected to UV-ozone treatment for 40 min to enhance surface reactivity prior to ECM coating.

Dissociated fibroblasts (1×10^7^ cells ml^−1^) were then introduced into the upper microchannel and allowed to adhere under static conditions for 2 h in a humidified 37 °C incubator with 5% CO_2_. After attachment, culture medium was perfused into the upper and/or lower microchannels at a constant flow rate of 20 μl h^−1^. To recapitulate peristalsis-like mechanical stimulation, cyclic deformation (10% strain, 0.15 Hz frequency) was applied via vacuum chambers interfaced with a computer-controlled tension system (FX-6000 Tension System with FlexLink Controller; Flexcell International Corporation). The negative pressure cycles deformed the elastic PDMS membrane, thereby transmitting dynamic strain to the fibroblast or epithelial cell layers cultured atop the membrane. To establish a dual-sided co-culture of iFib on both sides of a porous membrane, dissociated iFib cells (1×10^7^ cells ml^−1^) were first introduced into the upper microchannel and incubated under static conditions for 2 h at 37 °C in a humidified CO_2_ incubator. The device setup was then gently inverted, and the same iFib suspension was introduced into the opposing (now upper) microchannel, followed by a second 2-h static incubation to allow cell attachment. After ensuring iFib adhesion to both sides of the membrane, the device was returned to its upright orientation, and continuous perfusion of fibroblast culture medium was initiated at 20 μl h^−1^. For MMP inhibition experiments, the broad-spectrum MMP inhibitor GM6001 (25 μM; Calbiochem) was added to the fibroblast culture medium during microfluidic perfusion. For static control experiments, fibroblasts (1×10^7^ cells ml^−1^) were seeded either onto ECM-coated PDMS membranes placed in glass-bottom dishes or onto nanoporous membrane inserts and cultured under standard incubator conditions (37 °C, 5% CO_2_) for 5 days.

To perform epithelial–fibroblast co-cultures, the gut-on-a-chip device was preconditioned by perfusing an organoid culture medium through the upper microchannel and a fibroblast culture medium through the lower microchannel for 6 h at 20 μl h^−1^. Dissociated normal human intestinal epithelial organoid cells (∼1×10^7^ cells ml^−1^; passage number 5) were seeded into the ECM-coated upper channel and incubated under static conditions for 2 h to allow for attachment. Subsequently, dissociated fibroblasts were introduced into the lower channel (preconditioned with a fibroblast medium), and the device was inverted and incubated for an additional 2 h at 37 °C. The chip was then restored to its original orientation and continuous medium perfusion was resumed through both channels at 20 μl h^−1^.

### Isolation of mechanoadaptive intestinal fibroblasts

Normal intestinal fibroblasts (nFib) cultured under continuous apical shear flow (20 μl h^−1^) in a gut-on-a-chip exhibited extensive apoptotic detachment within 24 h, with a residual population forming multicellular aggregates enriched in fibrillar structures. On day 10, aggregates were enzymatically dissociated using 0.25% Trypsin-EDTA solution (Gibco; cat. no. 25200072) for 5 min at 37 °C. Dissociated cells were collected, filtered through a 70 μm strainer, and resuspended in a fibroblast culture medium. The isolated mechanoadaptive fibroblasts (nFib_MA_) were expanded under standard conditions (37 °C, 5% CO_2_) and, upon serial passaging, developed distinct morphological and phenotypic traits consistent with a myofibroblast-like state.

### Morphological analysis

Fibroblast morphology was monitored under various culture conditions, including T75 tissue culture flasks, nanoporous Transwell inserts, PDMS membranes, and gut-on-a-chip microdevices. Phase-contrast images were acquired using an inverted microscope (DMi1; Leica Microsystems) equipped with 10× (NA 0.22) and 20× (NA 0.30) objectives, a digital camera (MC120 HD; Leica), and Leica Application Suite software (LAS v4.12; Leica). For each condition and time point, images were captured from over 10 random fields of view across at least two independent biological replicates. Representative images were selected for figure presentation.

For immunofluorescence staining, cells were fixed with 4% (w/v) paraformaldehyde (Thermo Fisher Scientific) for 30 min, permeabilized with 0.3% (v/v) Triton X-100 (Millipore Sigma) for 15 min, and blocked with 2% (w/v) bovine serum albumin (BSA; Millipore Sigma) in phosphate-buffered saline (PBS; Gibco) for 1 h at room temperature. All steps included PBS washes between treatments. Primary antibodies targeting α-smooth muscle actin (α-SMA; mouse, Abcam, cat. no. ab7817), collagen I (COL I; rabbit, Abcam, cat. no. ab90395), and vimentin (mouse, Thermo Fisher, cat. no. MA5-11883) were applied for 1 h at room temperature. Samples were then incubated with Alexa Fluor-conjugated secondary antibodies (Alexa Fluor 555 goat anti-rabbit, Abcam, cat. no. ab150078; Alexa Fluor 488 donkey anti-mouse, Abcam, cat. no. ab150105) for 1 h in the dark. Nuclei and F-actin were counterstained using 4′,6-diamidino-2-phenylindole dihydrochloride (DAPI, 1 μg ml^−1^; Thermo Fisher, cat. no. 62248) and CruzFluor 647-conjugated phalloidin (1:500; Santa Cruz Biotechnology, cat. no. sc-363797). Samples were mounted using Fluoromount medium (Millipore Sigma, cat. no. F6182).

Confocal imaging was performed using a Leica TCS SP8 microscope (Leica Microsystems) equipped with a 25× water immersion objective (NA 0.95), excitation lasers (405 nm, 488 nm, 561 nm, and 633 nm), and a hybrid detector (HyD). Single-plane and Z-stack images were acquired and analyzed using Leica LAS X software. Orthogonal projections from Z-stack images were generated for high-resolution vertical cross-sectional analysis. Fluorescence intensities were quantified from three randomly selected areas per condition, and cell layer thickness was measured from 10 cross-sectional views using ImageJ (NIH).

For live-cell imaging, normal fibroblasts were labeled using CellTracker dye (Invitrogen; cat. no. 7025) according to the manufacturer’s instructions. Briefly, cells were harvested by trypsinization, resuspended in CellTracker working solution (final concentration, 10 μM), and incubated at 37 °C for 30 min. Labeled cells were then washed twice with PBS, resuspended in fibroblast culture medium at a concentration of approximately 1×10^7^ cells ml^−1^, and introduced into the upper microchannel of a gut-on-a-chip device. Following cell attachment under static conditions, fluorescence imaging was performed intermittently using an EVOS fluorescence microscope (EVOS M5000; Invitrogen) equipped with a 10× (NA 0.3) objective and an integrated live-cell monitoring system.

For scanning electron microscopy (SEM) analysis, cells were fixed with 2% (w/v) paraformaldehyde (Electron Microscopy Sciences, cat. no. 15710) and 2.5% (w/v) glutaraldehyde (Electron Microscopy Sciences, cat. no. 16320) in PBS for 30 min at room temperature, followed by post-fixation with 1% (w/v) osmium tetroxide (Electron Microscopy Sciences, cat. no. 19150) in 0.1 M sodium cacodylate buffer (Electron Microscopy Sciences, cat. no. 11650) for 60 min in a fume hood. Samples were extensively rinsed with PBS, dehydrated through graded ethanol solution (70%, 95%, and 100%, v/v; 10 min each), treated with hexamethyldisilazane (HMDS; Electron Microscopy Sciences, cat. no. 16700) for 10 min, and dried overnight in a vacuum desiccator containing anhydrous desiccant (Drierite, 8 mesh; cat. no. 23005). Dried specimens were mounted on aluminum stubs with conductive carbon adhesive tape, sputter-coated with ∼10 nm layer of gold (Electron Microscopy Sciences), and imaged using a Zeiss SIGMA VP Scanning Electron Microscope (Carl Zeiss Inc.).

The aspect ratio of normal and inflammation-associated fibroblasts was quantified using fluorescence images visualizing F-actin and nuclei. For each cell, the longest cellular axis (length, *L*) and the shortest perpendicular axis (width, *W*) were measured using ImageJ software. The aspect ratio (AR) was calculated as the ratio of length to width (AR = *L*/*W*), providing a quantitative measure of cell elongation. For each experimental condition, at least 10 cells were analyzed to obtain a representative average aspect ratio.

Fibroblast alignment in response to fluid shear stress or cyclic strain was assessed by measuring the percentage of cells oriented along the principal strain axis and analyzing their orientational angles relative to this axis. The degree of alignment was determined by calculating the percentage of cells oriented within a defined angular range relative to the direction of applied mechanical force, providing a quantitative measure of fibroblast alignment^67^. Alignment analysis was performed using the Directionality plugin in ImageJ software, which quantifies pixel orientation distributions from fluorescence images and generates histograms representing the predominant cellular alignment.

### Mechanobiological characterization

Mechanical properties of fibroblasts, cell aggregates, and culture substrates were quantified using a high-performance atomic force microscope (AFM; MFP-3D-Bio AFM, Oxford Instruments). Tipless cantilevers (ARROW-TL1Au, Nanoworld; nominal spring constant, 0.03 N·m^−1^) were modified by attaching 5-μm polystyrene beads (Polysciences, Inc.), as previously described^68^. For live-cell indentation assays, force-distance curves were collected from at least 25 randomly selected fibroblasts per condition, using a constant scan rate of 0.15 Hz and trigger setpoint of 0.3 V. Young’s modulus was calculated by fitting force-indentation data (indentation depth ∼500 nm) to the Hertzian contact model^69^. Nanoindentation assays were similarly conducted on fibroblasts aggregates harvested at day 4 and day 10 of culture. For AFM topography, cells and aggregates were gently fixed with 0.1% (w/v) paraformaldehyde for 2 min at room temperature, rinsed with distilled water, and imaged at 0.5 Hz scan rate, setpoint of 0.1 V, and integral gain of 10 over a 90×90 µm^2^ scan area.

### Cellular assessment

Oxidative stress and apoptosis were simultaneously evaluated using CellROX Orange Reagent (Thermo Fisher Scientific, cat. no. C10444) and BioTracker NucView 405 Blue Caspase-3 Dye (MilliporeSigma, cat. no. SCT-104), respectively. Fibroblasts cultured in a gut-on-a-chip were incubated with a mixture of both dyes (5 μM each) perfused into the apical and basal microchannels at 20 μl h^−1^ in a 5% CO_2_ incubator at 37 °C. After 30 min reaction, cells were rinsed with PBS and imaged by live-cell confocal microscopy. Reactive oxygen species (ROS) and caspase-3 activity were detected at excitation/emission wavelengths of 555/565 nm and 405/450 nm, respectively. For quantitative analysis, fluorescence intensities were measured from at least three randomly selected fields per condition using ImageJ software. All experiments were performed in technical duplicates and repeated in at least three independent biological replicates.

Fibroblast proliferation was assessed using the Click-iT EdU Alexa Fluor 488 Imaging Kit (Invitrogen, cat. no. 10337). Fibroblasts cultured under flow alone (fluidic) or combined flow and mechanical strain (dynamic) conditions in the gut-on-a-chip were incubated with 50 μM EdU in fibroblast medium for 3 h at 37 °C. After labeling, cells were fixed with 3.7% (w/v) PFA for 20 min, washed with 3% (w/v) BSA in PBS, permeabilized with 0.5% (v/v) Triton X-100 for 30 min, and stained with the Click-iT reaction cocktail for 30 min in the dark. Nuclei were counterstained with DAPI (1 μg ml^−1^). EdU-positive cells and total nuclei were imaged by confocal microscopy, and the percentage of proliferative cells was calculated as EdU+/total nuclei across at least three randomly selected fields per condition.

### Gene expression analysis

Total RNA was extracted from fibroblasts using the RNeasy Mini Kit (Qiagen) following the manufacturer’s protocol. RNA purity and concentration were assessed with a NanoDrop spectrophotometer (Thermo Fisher Scientific). Complementary DNA (cDNA) was synthesized from isolated RNA using the RT^2^ First Strand Kit (Qiagen; cat. no. 330404). Gene expression profiling in Fig. 3 was performed using the TaqMan Array Human Extracellular Matrix & Adhesion Molecules Panel (Thermo Fisher Scientific; cat. no. 4414133). Real-time PCR was conducted with the TaqMan Gene Expression Master Mix (Applied Biosystems; cat. no. 4369514) on a QuantStudio 3 Real-Time PCR System (Applied Biosystems). Relative gene expression levels were determined using the 2-ΔCT method, with normalization to HPRT1 as the housekeeping gene^70^. All reactions were performed in technical duplicates and averaged across at least three independent biological replicates.

### Protein quantification

The concentrations of TGF-β, IL-11, and COL1A1 were measured using DuoSet ELISA Kits (TGF-β, R&D Systems, cat. no. DY240-95; IL-11, DY218; COL1A1, DY6220-05) according to the manufacturer’s instructions. Standards and samples were added to 96-well plates pre-coated with capture antibodies, followed by sequential incubation with biotinylated detection antibodies and streptavidin-horseradish peroxidase. Signal was developed with a chromogenic substrate (3,3’,5,5’-tetramethylbenzidine, TMB), and absorbance was measured at 450 nm using a microplate reader (Cytation 5 Multi-Mode Reader; BioTek Instruments). Antigen concentrations were calculated from standard curves. All samples were run in technical duplicates and averaged across at least three independent experiments.

### Measurement of matrix metalloproteinase activity

Extracellular pan-MMP activity was quantified using the MMP Activity Assay Kit (Abcam; cat. no. ab112146) that covers various MMPs including MMP1, 2, 3, 7, 8, 9, 10, 12, 13, and 14^71^. Cell culture supernatant was collected, centrifuged at 1,000× *g* for 10 min, and mixed 1:1 (v/v) with 2 mM 4-aminophenylmercuric acetate (APMA) to activate latent MMPs, followed by incubation at 37 °C for 15 min. After addition of the fluorogenic pan-MMP substrate, fluorescence was measured every 10 min for 90 min at 37 °C using a microplate reader (BioTek Instruments) with excitation/emission settings of 490/525 nm. Each condition was assayed in technical duplicates and averaged across at least three independent biological replicates.

### Flow Cytometry

Fibroblasts were harvested via trypsinization, washed with 1% (w/v) BSA stain buffer (BD Biosciences), blocked with 5 μL Fc block (TruStain FcX, BioLegend; cat. no. 422301), then incubated on ice for 15 min. Cells were then fixed with BD Cytofix fixation buffer (BD Biosciences; cat. no. 554655) for 20 min on ice and permeabilized with BD Perm/Wash Buffer (BD Biosciences; cat. no. 554723). Staining was performed for 30 min at 4 °C in the dark using fluorophore-conjugated antibodies: anti-CD90-Brilliant Violet 421 (BioLegend; cat. no. 328114), anti-vimentin-Alexa Fluor 488 (BD Biosciences; cat. no. 562338), anti-α-SMA-APC (R&D Systems; cat. no. IC1420A), and anti-fibronectin-PE (R&D Systems; cat. no. IC1918P). After staining, cells were washed, resuspended in buffer, and analyzed on a Sony ID7000 Spectral Cell Analyzer (Sony Biotechnology). Data were processed using FlowJo software (Version 10; BD Biosciences). Fluorescence-minus-one (FMO) controls and isotype controls were used to establish gating strategies. Marker expression was quantified as the percentage of positive cells.

### Computational simulation

A two-dimensional computational model of laminar flow for an incompressible fluid was developed using COMSOL Multiphysics 5.4. The simulated domain included fluidic channels with heights of 500 μm (upper) and 200 μm (lower), separated by a porous membrane (10 μm pore diameter, 25 μm center-to-center spacing) representing the cellular interface. To model low Reynolds number conditions, inertial terms were neglected from the Navier-Stokes equations. Stationary studies were performed to evaluate flow velocity (μm s^−1^) and shear stress (μPa) distributions under four configurations representing epithelial (30 μm height) and fibroblast (2.5 μm height) layers, based on experimental observations. Cells were modeled as no-slip wall boundaries. Boundary conditions included fully developed laminar flow at the inlets (100 μl h^−1^ in the top channel; no flow in the bottom channel) and 0 Pa pressure at the outlets with backflow suppression. Velocity fields (“spf.u”, “spf.u.x”, and “spf.u.y”) were visualized using surface and vector plots. Shear stress was calculated by multiplying the shear rate (“spf.sr”) with dynamic viscosity (“spf.mu”), approximated as that of water (8.9×10^−4^ Pa·s). Shear stress values were exported as spreadsheets and filtered by x- and y-coordinates to extract data from regions corresponding to the cellular surfaces.

### Statistical analysis

All statistical analyses were performed using GraphPad Prism (version 10; GraphPad Software Inc.). Data are presented as the mean±standard error of the mean (SEM), unless otherwise specified. For comparisons between two groups, either two-tailed unpaired or paired Student’s t-tests were applied, depending on the experimental design. Nonparametric comparisons involving multiple groups were evaluated using the Kruskal-Wallis test followed by Dunn’s post hoc analysis, while nonparametric comparisons between two independent groups employed the Mann-Whitney U test. The ECM contraction assays were quantified by calculating the area under the curve (AUC), with statistical differences assessed using a paired t-test. A *p*-value less than 0.05 was considered statistically significant.

## Acknowledgements

This work was supported in part by the Kenneth Rainin Foundation Innovator Award (to H.J.K. & O.A.), the Crohn’s and Colitis Foundation of America Senior Research Award (to H.J.K.), Cleveland Clinic VeloSano Pilot Grants (to H.J.K.), Cleveland Clinic Catalyst SPARK Award (to H.J.K.), the Bio-industrial Technology Development Program from the Ministry of Trade, Industry & Energy Korea (20018770; to H.J.K.), Clinical and Translational Science Collaborative of Northern Ohio funded by the NIH NCATS, Clinical and Translational Science Award (UM1TR004528; to H.J.K.), and National Science Foundation (1927602, 1337859, and 2042116; to C.R.K.). Figures are partially created in BioRender.com.

## Author Contributions

S.M. and H.J.K. designed the study, performed experiments, and wrote the manuscript. N.T. performed and analyzed computational simulation. Y.C.S. and O.A. analyzed the data. E.G.E. and C.R.K. contributed to AFM experiment and analysis. All authors reviewed and approved the final version of the paper.

