## Supplementary Information for "Mechanostimulatory cues determine intestinal fibroblast fate and profibrotic remodeling in a physiodynamic human gut-on-a-chip"

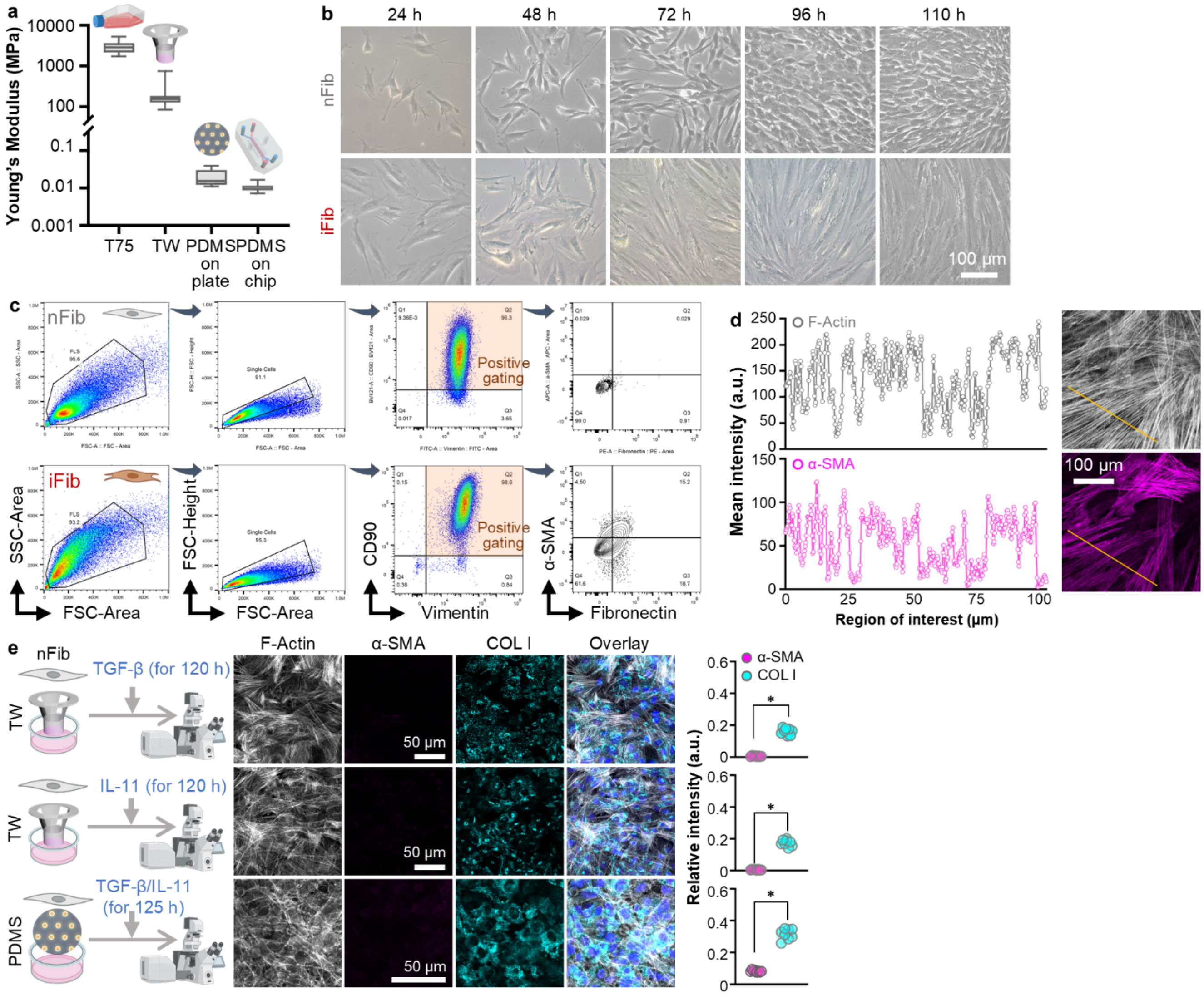


**Supplementary Fig. 1. Characterization of normal fibroblasts (nFib) and inflammation-associated fibroblasts (iFib) across various culture substrates. a**, Young's modulus measurements of different culture substrates, including a T75 flask, a nanoporous polyester membrane in a Transwell (TW), a PDMS membrane (10:1 base polymer:curing agent ratio), and a PDMS membrane fabricated within the gut-on-a-chip device (*n*=65 per condition). **b**, Representative phase-contrast images showing morphological profiles of nFib and iFib over time. **c**, Gating strategy for flow cytometry analysis of nFib and iFib cultured in T75 flasks. Cells were first gated based on forward scatter (FSC) and side scatter (SSC) to exclude debris and doublets, followed by gating for CD90 and vimentin expression (orange rectangular gates). Myofibroblast markers, α-SMA and fibronectin, were assessed within the positively gated populations. **d**, Mean intensity profile from line scan analysis of immunofluorescence confocal images of iFib cultured on a Transwell insert (related to Fig. 1e, lower panel). Orange lines indicate the regions of interest. F-actin (gray); α-SMA (magenta). **e**, Effect of treatment with profibrotic factors, TGF-β (10 ng ml$^{-1}$) and/or IL-11 (10 ng ml$^{-1}$), on nFib cultured on TW or PDMS membranes under static conditions for 120 h. The schematic (left) depicts the experimental setup. Representative immunofluorescence micrographs show F-actin (gray), α-SMA (magenta), and COL I (cyan) expression, along with overlay images. Quantification of relative fluorescence intensities is shown (right panel) ($n=4$). *$p<0.001$.

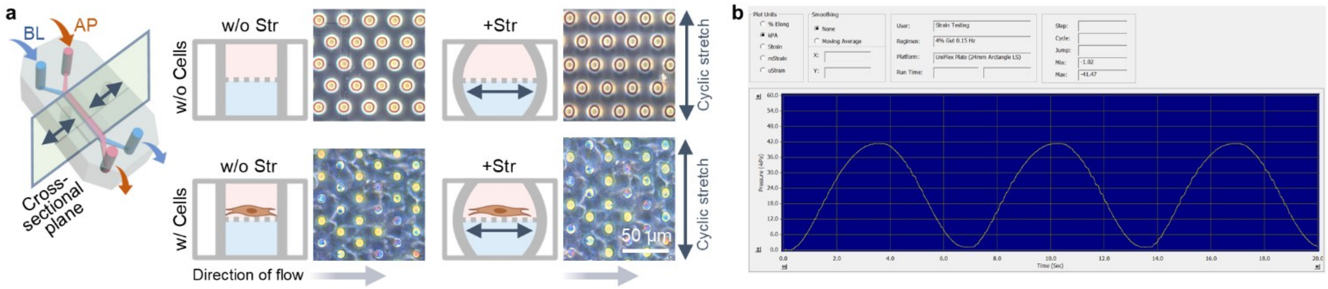


**Supplementary Fig. 2. Operational mechanism of cyclic stretch in a gut-on-a-chip microsystem. a**, Schematic illustrating the application of biaxial cyclic stretch and unidirectional flow to cells cultured within the gut-on-a-chip microfluidic device. The top panels show a cross-sectional view of the device without cells under static conditions (w/o Str) and during cyclic stretch (+Str). The bottom panels depict corresponding conditions with cells adherent to the porous membrane substrate. Phase contrast micrographs of cell-free PDMS membranes and membranes with adherent cells are shown. Grey arrows indicate the direction of culture medium flow; blue double-headed arrows represent the direction of cyclic mechanical stretch. **b**, Sinusoidal waveform depicting the cyclic stretch profile applied to the microfluidic device via a vacuum-driven pneumatic regulator (Flexcell Tension System).

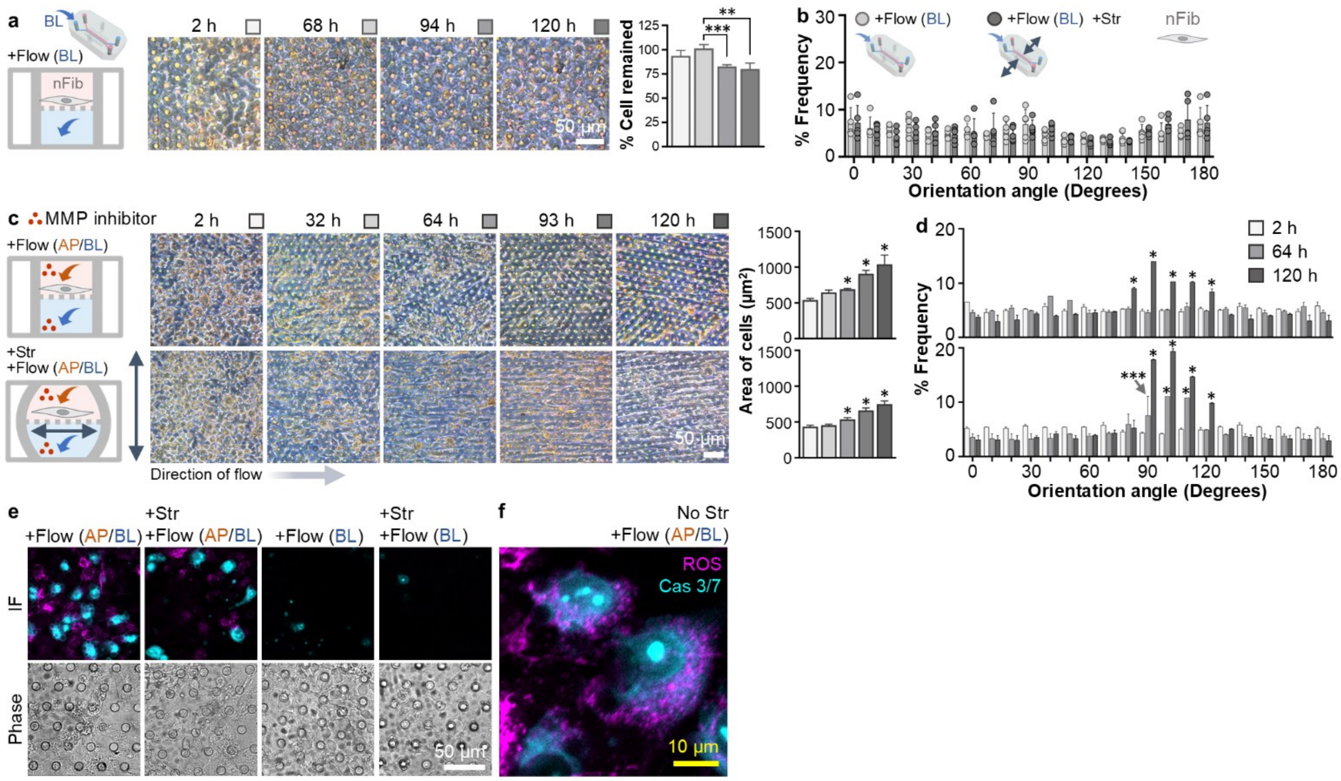


**Supplementary Fig. 3. Cellular and morphological assessment of nFib under various biomechanical conditions or MMP inhibition. a**, Time-lapse phase-contrast images showing nFib morphology over time under pseudostatic basolateral flow (+Flow BL). The right panel shows quantification of the percentage of cells remaining at each time point (*n*=3). **b**, Polar histograms depicting the frequency distribution of orientation angles for nFib cultured in a gut-on-a-chip under basolateral flow alone (+Flow BL) or combined basolateral flow and cyclic mechanical strain (+Flow BL, +Str), analyzed at 120 h using ImageJ software. **c**, Representative phase-contrast micrographs of nFib treated with MMP inhibitor (GM6001, 25 μM) under dual flow conditions (+Flow AP/BL) with or without cyclic strain (+Str). Grey arrows indicate the direction of medium flow. Quantification of cell density over time is shown on the right (*n*=10), with statistical comparisons made relative to cell numbers at 2 h. **d**, Polar histograms showing the orientation angle distribution of nFib cells cultured in a gut-on-a-chip treated with MMP inhibitor (related to **c**) (*n*=5). **e**, Representative fluorescence micrographs (upper panels) showing ROS (magenta) and caspase-3/7 (cyan) activity, along with corresponding phase-contrast images (lower panels), for nFib cultured under dual flow (+Flow AP/BL) or pseudostatic basolateral flow (+Flow BL) with or without cyclic stretch (+Str) at 120 h. **f**, High-magnification fluorescence micrographs highlighting ROS and caspase-3/7 signals in nFib challenged with dual flow without cyclic strain at 120 h. *$p<0.001$, **$p<0.01$, ***$p<0.05$.

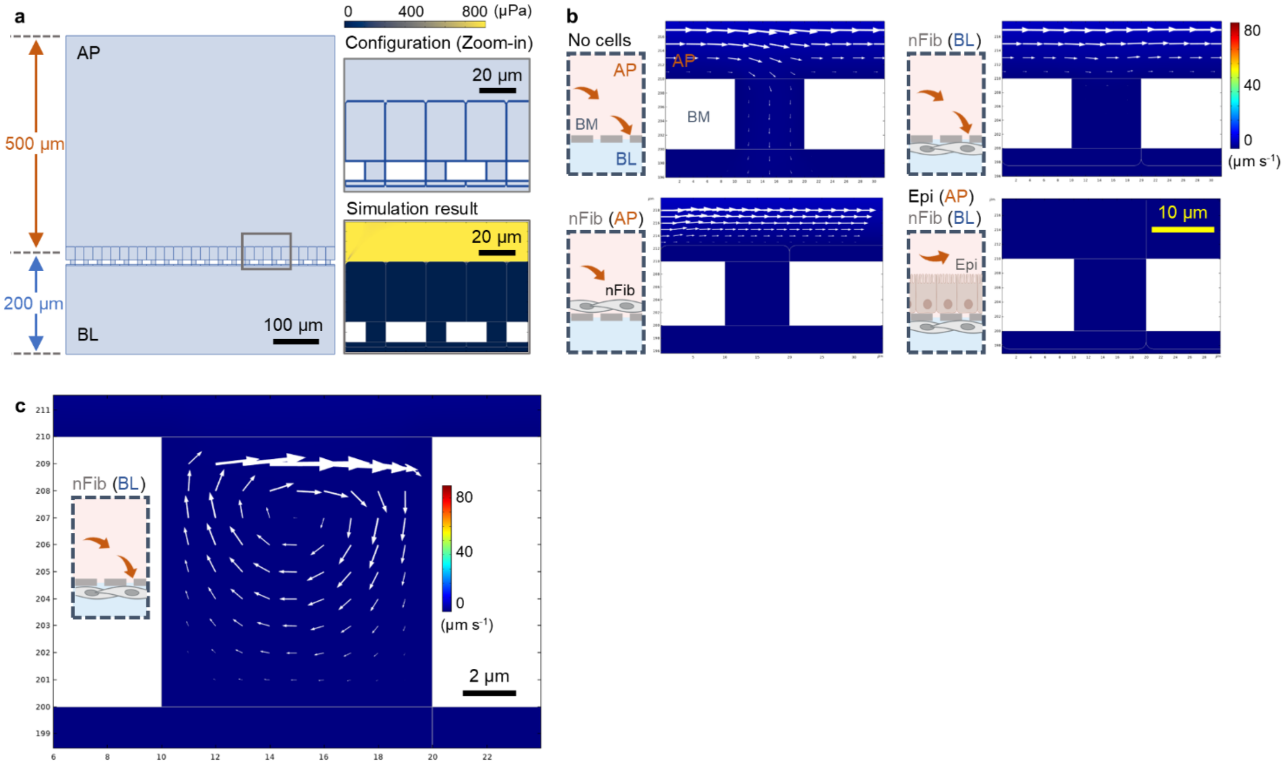


**Supplementary Fig. 4. Configuration of computational simulations and flow velocity profiles under various culture conditions. a**, a, Schematic illustrations showing a vertical section of the gut-on-a-chip microchannels (left) and a magnified view of the epithelial-nFib interface (top right). A representative simulation result of flow velocity is shown (bottom right). **b**, Computationally estimated flow velocity profiles for different culture configurations. The size and thickness of white arrows indicate the magnitude of velocity. Configurations include: no cells, nFib seeded on the apical side (nFib AP), nFib seeded on the basolateral side (nFib BL), and epithelial cells on the apical side with nFib on the basolateral side (Epi AP, nFib BL). **c**, Zoomed-in view of the two-dimensional velocity profile in the condition where nFib cells were cultured underneath the porous membrane. Rotational eddies formed under steady-state flow are visualized by local variations in flow velocity.

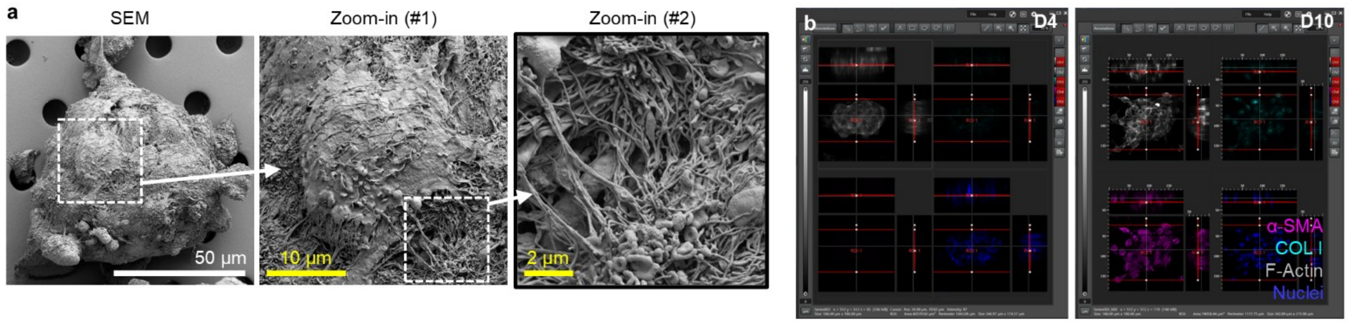


**Supplementary Fig. 5. Morphological characterization of D10 nFib aggregates. a**, Scanning electron microscopy (SEM) images of a D10 nFib aggregate showing the surface morphology and internal extracellular matrix (ECM) structure. Dashed white squares indicate the region of serial zoom-in views (#1 and #2) that highlight the complex fibrillar network. **b**, Configurational regions of interest (ROIs) for confocal micrographs of D4 and D10 aggregates (related to Fig. 5d and 5e), illustrating the spatial distribution of fibrotic markers, including α-SMA and COL I, counterstained with F-actin and nuclei.

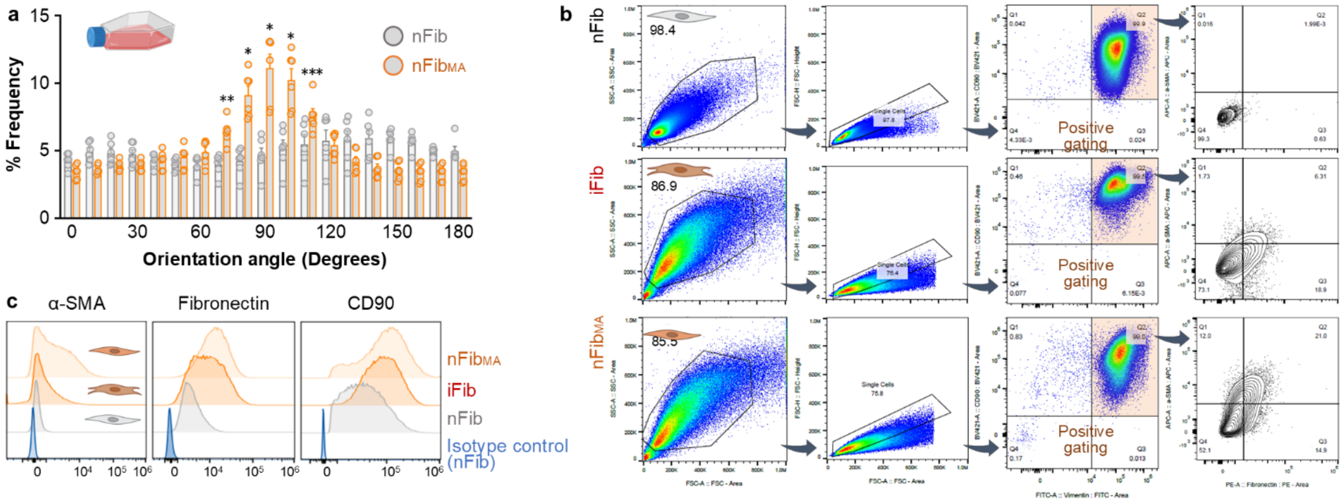


**Supplementary Fig. 6. Characterization of isolated mechanoadaptive fibroblasts ($nFib_{MA}$). a**, Polar histograms illustrating the orientation angle distribution of nFib and $nFib_{MA}$ cultured in T75 flasks for 96 h, analyzed using ImageJ software (*n*=6). **b**, Gating strategy used for flow cytometry analysis of nFib, iFib, and $nFib_{MA}$ cultured in T75 flasks (96 h). Cells were initially gated by forward scatter (FSC) and side scatter (SSC) to exclude debris and doublets, followed by gating for CD90 and vimentin expression (orange rectangular gates). Expression of myofibroblast markers α-smooth muscle actin (α-SMA) and fibronectin was assessed within these positively gated populations. **c**, Flow cytometry histograms comparing expression profiles of fibrosis-associated markers (α-SMA, fibronectin, and CD90) among nFib, iFib, and $nFib_{MA}$ cultured in T75 flasks.
